# Using sequence-to-function models to interpret archaic hominin introgression

**DOI:** 10.64898/2026.08.31.748430

**Authors:** Maddy Comerford, Isabela Alvim, Timothy M. Johanson, Danat Yermakovich, Christopher Kinipi, Matthew Leavesley, François-Xavier Ricaut, Michael Dannemann, Murray P Cox, Rhys Allan, Nicolas Brucato, Irene Gallego Romero

## Abstract

Understanding the functional impact of archaic hominin introgression remains challenging due to the poor representation of global introgression in publicly available genomics resources. Sequence-to-function models can predict the effects of any possible variant in the human genome and may fill this gap. Here, we used AlphaGenome to predict the effects of 144,139 introgressed SNPs segregating in present-day individuals of Papuan genetic ancestry. AlphaGenome’s chromatin accessibility predictions recapitulate experimentally observed effects, but gene expression performs no better than chance. Predictions correlate more strongly with an independent reporter assay of single-variant activity than with the same variants’ effects in live cells, indicating that AlphaGenome captures the regulatory potential of individual variants more reliably. Predictions carry tissue specificity, allowing us to predict specific tissues potentially impacted by introgressed haplotypes. We identify genes, including JAK1 and TAB2, that are associated with haplotypes that contain an excess of variants predicted by AlphaGenome to have large impacts on chromatin accessibility. Finally, we highlight the challenges and limitations associated with using sequence-to-function models for introgressed variant effect prediction, and show that while AlphaGenome’s chromatin accessibility predictions can aid in prioritising candidate functional regions, expression predictions and the assignment of variants to target genes remain as open challenges.

## Introduction

Introgression from Neanderthals and Denisovans is a defining feature of the genomes of present-day human populations. As a result of multiple encounters between modern humans and these archaic hominins, the genomes of present-day non-African individuals is made up of up to 1-3% Neanderthal DNA [1, 2, 3], while some individuals, particularly those of Papuan or Near Oceanic genetic ancestry, owe up to an additional 4% of their DNA to Denisovans [4, 5, 6]. Understanding the phenotypic consequences of archaic hominin introgression remains an active area of research, with results suggesting consistent contributions from both Neanderthals and Denisovans to immune function and metabolism [7, 8], alongside more limited contributions to genes impacting a suite of phenotypes from skin pigmentation [9] to hypoxia tolerance [10]. However, there are multiple challenges to studying archaic introgression. Introgressed DNA largely persists in non-coding regions [11, 12, 13], where interpreting variant effect can be difficult. The size of current functional genomics resources, which rarely exceed 1,000 individuals, makes resolving the impact of introgressed variants challenging, as these generally segregate at lower allele frequencies. Additionally, the bias towards individuals carrying Neanderthal introgression in publicly available databases limits the study of Denisovan introgression.

Some of these limitations are easier to overcome than others. For instance, the recent eQTLGen II consortium brings together gene expression profiles in whole blood from over 43,000 samples, enabling the identification of functional cis and trans regulatory variants at unprecedented resolution [14]. This enables the fine-grained exploration of the impact of Neanderthal introgression on gene expression levels in blood, even when they segregate at low frequencies. However, donors of non-European genetic ancestries remain poorly represented, and account for approximately 10% of samples in eQTLGen II, restricting the ability to explore Denisovan introgression. Likewise, while whole blood is relatively straightforward to collect and profile, representation of global genetic diversity in datasets that can only be collected through more invasive or post-mortem sampling is more limited [15], making it more challenging to resolve the functional consequences of Denisovan introgression across tissues.

“Sequence-to-function” (S2F) models are a growing family of deep learning models that can predict multiple modalities of functional genomic data from DNA sequence alone [16, 17]. Trained on publicly available data, these models aim to learn underlying sequence features that determine function, allowing them to theoretically predict the effect of any possible variant in the human genome. Existing models can be either specialised in the prediction of one or a limited number of closely related data modalities, such as ChromBPNet for chromatin state [18] or SpliceAI for splicing [19], or generalist predictors that operate across a broad range of data modalities, such as Enformer, Borzoi or AlphaGenome [20, 21, 22]. Single-modality models have already been used to examine alternative splicing [23] and higher order chromatin structure [24] in the context of archaic hominin introgression. However, the accuracy of these models is known to vary across modalities, and a dearth of experimental data limits validation. Model performance appears to be more accurate when predicting “local” effects, such as the impact of a variant on chromatin accessibility in its immediate surroundings, than when predicting across large genetic distances, such as when predicting the effect of cis-eQTLs, [22, 25]. These and other factors have limited the application of these methods to making individual-level expression predictions [26, 27], with state of the art tools frequently performing no better than linear models with orders of magnitude fewer parameters such as prediXcan [28].

Here, we generate AlphaGenome predictions for over 144,000 introgressed SNPs segregating in present-day Papuan populations, and assess the utility of AlphaGenome in interpreting introgressed variant effect. Crucially, we make use of both existing and newly generated functional genomics data to validate the accuracy of these predictions, allowing us to establish boundaries for the use of AlphaGenome’s predictions. We outline challenges that limit the application of AlphaGenome predictions for understanding archaic hominin introgression, but show that these predictions both capture known biological trends and extend current knowledge. Thus, AlphaGenome holds promise as a tool to examine the functional consequences of introgression from archaic hominins into present-day individuals.

## Materials and Methods

### Ethics approvals and human samples

The 11 Papuan LCLs used in this study were established from healthy adult donors sampled at the University of Papua New Guinea, in Port Moresby, Papua New Guinea between 2018 and 2019 following the procotol outlined in [29]. Collection of blood was approved by the Medical Research Advisory Committee of the National Department of Health of the Government of Papua New Guinea (permit number MRAC 16.21), by the University of Melbourne’s Human Research Ethics committee (approvals 1851585.1 and 26981), and by the French Ethics Committee (Committees of Protection of Persons 25/21_3, n*^◦^*SI:21.01.21.42754). Permission to conduct research in Papua New Guinea was granted by the National Research Institute of Papua New Guinea (permit 99902292358), with full support from the School of Humanities and Social Sciences at the University of Papua New Guinea. All individuals gave their full informed written consent to participate in this study.

### Genome sequencing and introgression calling

All genomes used in this study have been previously published [30, 31]. Briefly, DNA was obtained from saliva samples and sequencing libraries were prepared using the TruSeq DNA PCR-Free HT kit (Illumina, Inc., San Diego, California, USA). Sequencing was performed using a 150-bp paired-end protocol on the Illumina HiSeq X5 sequencer (Illumina, Inc., San Diego, California, USA). All data are deposited and available on the European Genome-Phenome Archive under accession numbers EGAD00001010142, EGAD00001010143, EGAD50000000050, and EGAS00001005393.

We used a previously established workflow for reconstructing and annotating archaic introgressed haplotypes [32], which consists of the following steps: i) identifying plausible introgressed archaic SNPs that are absent in YRI population from 1000 Genomes project but shared between target individuals and archaic hominins; ii) group them into haplotypes based on linkage disequilibrium (LD); iii) test the haplotypes for incompatibility with incomplete lineage sorting; iv) compare the sequence similarity of inferred archaic haplotypes to Neanderthal and Denisovan genomes to define the source of introgression. We applied the published version of the method to 249 whole-genome sequences from individuals in Papua New Guinea described above; this included the 11 donors with established LCL lines used in this study. We removed haplotypes containing fewer than five SNPs or those where incomplete lineage sorting could not be ruled out. We assigned aSNPs to their nearest TSS using gene annotations from Ensembl v110 (hg38). Genomic annotations for aSNPs were generated with HOMER [33].

### Cell culture

The 11 Papuan LCLs were cultured in suspension in RPMI with L-glutamine (Sigma, for RNA-seq; Gibco, for ATAC-seq) supplemented with 1% NEAA (Gibco), 1% Glutamax, and 10% FBS (Gibco, for ATAC-seq) or 15% FBS (Scientifix, for RNA-seq). Nine LCLs were used for RNA-seq data generation; for ATAC-seq data generation we cultured an additional 2 lines, for a total of 11 LCLs.

### RNA-seq data generation

2 *×* 10^6^ LCLs were harvested by centrifugation and RNA extracted using the Qiagen’s RNeasy Mini Kit following manufacturer’s protocols. Directional mRNA library preparation (poly A enrichment) and 150bp paired end sequencing on Illumina NovaSeq X Plus were performed by Novogene to an average depth of 30 million read pairs. Read quality was assessed with FastQC v0.12.1 [34]. Sequencing adapters were removed with trimmomatic v0.39 [35] and trimmed reads were passed to STAR v2.7.11b [36] for two-pass alignment to the human genome (hg38, Ensembl v110). Mapped reads were passed to featureCounts v2.0.8 [37] for read quantification. Genes with log_2_CPM *≥* 1 in at least half of the samples were retained for downstream analyses.

### ATAC-seq data generation

ATAC-seq library preparation was performed using an optimised version of the Omni-ATAC protocol [38]. 50,000 LCLs were pelleted at 500g at 4*^◦^*C for 5 minutes and resuspended in 50*µ*l cold ATAC-resuspension buffer (RSB, containing 10mM Tris-HCl pH 7.4, 10mM NaCl, 3mM MgCl2, 0.1% Igepal, 0.1% Tween-20 and 0.01% Digitonin). This reaction was incubated on ice for 3 minutes and then mixed with 1ml cold ATAC-RSB (containing 0.1% Tween-20 but not Igepal or Digitonin). Nuclei were pelleted at 500g for 10 minutes at 4*^◦^*C and resuspended in 50*µ*l transposition mixture (25*µ*l 2x RD buffer, 2.5*µ*l transposase, 16.5*µ*l PBS, 0.1*µ*l 5% Digitonin, 0.5*µ*l 10% Tween-20 and 5*µ*l water) by pipetting up and down 6 times. This reaction was incubated at 37*^◦^*C for 30 minutes in a thermomixer with 1000RPM mixing, after which 180*µ*l of ATL buffer was added. Purification was performed with Qiagen’s Mini-purification kit, adding 20*µ*l Proteinase K, following manufacturer’s instructions and then eluting in 20*µ*l elution buffer. The purified product was amplified in an 5-cycle PCR reaction with NEBNext 2x MasterMix. Immediately after this amplification, qPCR was performed to determine the additional number of PCR cycles required. After running the additional PCR cycles, reactions were purified using NucleoMag NGS Clean-up and Size Select, and libraries quantified with TapeStation (D5000), selecting a region from approximately 100-3000 bp. Data was sequenced (65 bp, paired end) at the Walter and Eliza Hall Institute’s core facility, to an average depth of 76 million read pairs, using the Illumina NextSeq platform.

ATAC-seq data was processed with the ENCODE pipeline (v1.8.0) available at https://github.com/ENCODE-DCC/atac-seq-pipeline. Raw sequencing reads were preprocessed to remove low-quality bases and adapter sequences with TrimGalore [39]. Cleaned reads were aligned to the human reference genome (hg38) using BWA-MEM [40] and duplicate reads were removed using Picard *MarkDuplicates* [41]. Transposase insertion sites were identified by converting aligned reads into tagAlign using BEDTools [42]. Peaks were called using MACS2 [43] with ENCODE pipeline standard parameters. Peaks located in hg38 blacklist regions [44] were removed. We defined a set of consensus peaks by merging overlapping peaks (*≥* 1bp) and retaining only peaks that are supported by original peaks in at least half the samples. Reads mapping to consensus peaks were counted with the regionCounts() function [45] and converted to reads per kilobase per million mapped reads (RPKM).

### Estimating differential activity in RNA-seq and ATAC-seq

To determine whether genotype at an aSNP is associated with significant differences in the experimental RNA-seq and ATAC-seq datasets, we consider only aSNPs in which at least three individuals (of nine for RNA-seq, and of 11 for ATAC-seq) carry the introgressed allele (either as homozygotes or heterozygotes). Existing differential expression frameworks such as limma or DESeq2 borrow information genome-wide to stabilise the mean/variance relationship and control for overdispersion in RNA-seq data. However, because introgression is unevenly distributed across the genome, we cannot fit a single linear model to our data, making the use of these models impossible. As such, at each site we instead quantify differences between introgression and non-introgression carriers using Hedges’ g, a standardised mean difference that scales the shift in expression or accessibility between groups by the within-group variability and additionally controls for biases introduced by our small sample size.

### AlphaGenome predictions

We generated computation predictions of variant effects with AlphaGenome [22]. All AlphaGenome predictions were generated using the publicly available API and the *dna_model.score_variant* command. We used 1Mb surrounding the variant of interest as input, and generated RNA-seq and ATAC-seq predictions for a subset of available Biosamples by specifying the relevant cell ontologies (**Supplementary table 1**). We chose to generate predictions for Biosamples that are broadly representative of cell types and tissues previously described as relevant to archaic introgression [13]. We generated both baseline active scores that captures the absolute activity level of the SNP (ACTIVE scorer for both RNA-seq and ATAC-seq), and a log_2_ fold change score that reflects the difference in scores between alleles of a variant (for RNA-seq: GeneMaskLFCScorer, for ATAC-seq: CenterMaskScorer). For RNA-seq scores, we keep only scores for each aSNP-nearest gene pair, and consider only polyA-plus genes. For both modalities, we polarised log_2_ fold change score scores to reflect the difference between the introgressed and non-introgressed alleles.

We also compared AlphaGenome predictions to MPRA data from [46]. Variants in the MPRA data from [46] were classified as being simply Denisovan or Neanderthal, so we combine all Neanderthal variants into a single ancestry group for these analyses. Bootstrap confidence intervals for comparisons between the three data modalities (MPRA, AlphaGenome and experimental data were calculated from 10,000 resamples. The change in correlations was calculated as the difference in correlations within a modality between MPRA and in-vivo data. When performing PCA, and to control for a possible lack of independence in scores from aSNPs located in the same haplotype we randomly selected one aSNP per haplotype (n = 7,956 for ATAC-seq, 7,409 for RNA-seq), and repeated this process 500 times, aligning the results using Procrustes.

#### Raw and quantile scores

AlphaGenome predictions are reported as both raw scores and a normalised, empirically derived quantile score. Raw scores are not normalised between tracks or modalities, but can be informative for examining the absolute magnitude of effects within a track. Quantile scores are generated by comparison against a genome-wide background distribution of approximately 300,000 variants with AF *≥* 0.01 in at least one gnomAD v3 population; this set is held constant across all tissues and data modalities [22]. This means that quantile scores represent a variant’s rank within this invariant background variant distribution, which enables direct comparisons between tracks and scorers.

Quantile scores can be either signed or unsigned, depending on whether the raw score is itself directional. Thus, for the baseline activity (ACTIVE) predictions, which are unsigned, quantile scores range form 0 to 1, but for the differential activity predictions (GeneMaskLFCScorer and CenterMaskScorer), which are signed, they are bounded between -1 and 1, with the two extremes representing the most extreme negative (0th percentile) and positive (100th percentile) observations within each track. For these reasons, and following the AlphaGenome documentation, we perform comparisons between Biosamples and data modalities considering only quantile scores, but also consider raw scores when examining each Biosample individually. Differential activity scores are always reported relative to the introgressed allele, with a positive quantile indicating higher activity is associated with the introgressed allele than with the non-introgressed allele.

### Identifying haplotypes with excess signal

We considered two different ways to examine the clustering of *|q|* scores within a haplotype.

i. In the simpler approach, we compute the overall fraction of aSNPs with *|q|* exceeding a threshold for each Biosample; e.g. *|q| ≥* 0.9 in B cells = 0.105. We use this probability to determine whether a haplotype has an excess of aSNPs with a *|q|* score that exceeds the threshold under the binomial distribution, where p = the observed Biosample*×*threshold fraction, n = the number of aSNPs in a haplotype, and k = the observed number of aSNPs in the haplotype that exceed the threshold. We only perform this test on haplotypes with at least 5 scored aSNPs and at least one aSNP that exceeds the *|q|* threshold, and apply BH correction across all haplotypes*×*Biosample tests for each threshold. This approach effectively asks “conditioning on a haplotype containing at least one aSNP with *|q| ≥ threshold*, do we observe more aSNPs with *|q| ≥ threshold* than we expect?”
ii. The approach above does not control for the observed excess of signal near the TSS. To do this, we use a two-step procedure. First, we use logistic regression to calculate the probability that a given aSNP has a *|q|* score above a given threshold, fitting regressions independently for each Biosample and *|q|* threshold. In each case, we binarise the AlphaGenome *|q|* score into above or below threshold for that aSNP and Biosample and fit the following model:

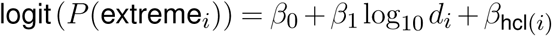

where log_10_d*_i_* is the log_10_ distance between the aSNP to its nearest TSS, and hcl is the aSNP’s annotation as given by HOMER [33], treated as an unordered factor with the following possible levels: intergenic (71,121 aSNPs), intron (66,140), promoter-TSS (1,802), TTS (1,586), 3’ UTR (1,466), exon (1,133), non-coding (751), 5’ UTR (140). Biosample/threshold combinations were skipped if there were fewer than 50 aSNPs above the threshold.

We then use these probabilities to generate a baseline expectation for how many aSNPs with *|q| ≥ threshold* we expect for a given haplotype. To do this, we take into account the fact that each aSNP has a different probability p*_i_* of exceeding the *|q|* threshold, making the expected number of extreme *|q|* aSNPs the sum of *n* independent Bernoulli trials with different success probabilities, where *n* is the number of aSNPs in a haplotype. For each haplotype we then compare the observed number of aSNPs with *|q| ≥ threshold* to this expectation under the Poisson-binomial distribution. As above, we only perform this test on haplotypes with at least 5 scored aSNPs and at least one aSNP that exceeds the *|q|* threshold, and apply BH correction across haplotypes independently for each Biosample*×*threshold combination. Significant haplotypes are those where the observed number of extreme *|q|* aSNPs is too high to be consistent with scores actually being independent.

We also used both approaches to identify haplotypes with a deficit of *|q| ≥ threshold* aSNPs. However, under a binomial distribution we are much better powered to detect an excess of signal than a deficit for haplotypes with a smaller number of SNPs; to control for this we limit deficit tests to haplotypes where the probability of observing zero aSNPs where *|q| ≥ threshold* is *<* 0.05. Under the Poisson-binomial model different haplotypes have different expectations, but here again we limit the testing to only haplotypes with power to detect a deficit. The number of tested haplotypes under each approach varies per Biosample and *|q|* threshold and is available as **Supplementary table 2** and **Supplementary table 3**. We considered the following thresholds for enrichment testing: *|q| >* 0, 7*|*0.8*|*0.9*|*0.95; distance to the nearest TSS *<* 1kb|5kb|10kb|20kb|50kb. No terms were significant at the 1kb threshold so it was ommited from figures and supplementary tables.

All analyses were performed in R 4.4.1 [47].

### Data access

Raw RNA-seq and ATAC-seq data have been deposited to EGA. Genotype data for all individuals can be accessed at EGA under accession numbers EGAD00001010142, EGAD00001010143, EGAD50000000050, and EGAS00001005393. Analysis code is available at https://gitlab.svi.edu.au/igr-lab/asnps_alphgen/.

## Results

### AlphaGenome can predict effects of archaic introgression on chromatin accessibility

To explore the relationship between AlphaGenome (AG) predictions and experimental observations of functional effects for archaic introgressed variants, we generated gene expression (RNA-seq) and chromatin accessibility (ATAC-seq) data from 11 lymphoblastoid cell lines (LCLs) established from unrelated donors sampled in Port Moresby, Papua New Guinea (see Methods). High coverage (30x) whole genome sequences were generated for all individuals as part of [31]; we used the set of 249 Papuan individuals in that study to identify archaic introgression in the 11 samples, totalling 144,139 alleles of confident Denisovan and Neanderthal ancestry, 67,170 of which were polymorphic in the 11 individuals with experimental data. We separated the Neanderthal introgressed SNPs (aSNPs) into two non-overlapping categories, “Neanderthal 1KG” and “Neanderthal PNG”, as in [32]. The “Neanderthal 1KG” label captures introgressed aSNPs residing on haplotypes that also segregate within mainland Eurasian individuals from the 1000 Genomes Project while “Neanderthal PNG” refers to haplotypes that have a closer match to one or more Neanderthal genomes than the Denisovan genome, but only segregate within Papuan individuals, suggesting a separate demographic history. In total, we labelled 52,705 aSNPs as Denisovan, 40,583 as Neanderthal 1KG, and 50,851 as Neanderthal PNG. We used AlphaGenome to predict the maximum impact of each aSNP on expression of nearby genes (RNA-seq) and chromatin accessibility (ATAC-seq) levels, as well as log_2_ fold changes between chromosomes carrying the introgressed and non-introgressed allele in GM12878, an LCL established from a female donor of European-like genetic ancestry. AlphaGenome RNA-seq predictions span all genes within an approximately 1MB stretch of genomic sequence centred on the tested aSNP. When aSNPs overlapped a protein coding gene, we retained the prediction for that gene; when aSNPs fell outside a gene, we retained the prediction for the gene with the nearest TSS, for a total of 6,094 genes with predictions across 129,004 aSNPs—because introgression is inherited in linkage blocks, the nearest gene is shared across many aSNPs, while aSNPs without predictions generally arise from there being no genes in the scored 1MB interval. ATAC-seq predictions are limited to a 501bp interval centred on the focal aSNP for all 144,139 aSNPs.

We first asked whether AlphaGenome’s baseline predictions of overall activity levels in GM12878 replicated observations made in LCLs from Papuan donors (Supplementary Figure 1). Ranking aSNPs by predicted accessibility recovered open chromatin well (AUROC = 0.956; Supplementary Figure 1A), with 79.4% of variants located within 250bp of an observed peak falling in the top decile of accessibility predictions and 92.4% in the top two (Supplementary Figure 1B). To perform the equivalent test for expression we ranked genes by their predicted RNA-seq coverage and asked whether they were deemed expressed in our experimental data, again observing good concordance (AUROC = 0.942; RNA-seq data is not available for 2 of the 11 cell lines). Because introgressed variants cluster on shared haplotypes and thus frequently have the same nearest gene, we repeated the accessibility test after thinning to one variant per 100 kb; the result is unchanged (AUROC = 0.952). Absolute measurements and predictions of activity also correlated well. For expression, Spearman *ρ* = 0.80 between predicted RNA-seq coverage and mean observed log_2_ CPM (n = 3,454 genes, Supplementary Figure 1C); for accessibility, *ρ* = 0.71 between predicted ATAC signal and mean peak RPKM (n = 8,945 variant–peak pairs from 8,736 variants, Supplementary Figure 1D).

This suggested that we could use these results to identify introgressed variants that directly impact gene expression levels or chromatin accessibility. For these analyses, we consider only aSNPs in which at least three individuals are either heterozygous or homozygous for the both introgressed and the non-introgressed alleles, allowing us to calculate variance for both genotype classes (introgression carriers and non-carriers). When considering RNA-seq, this resulted in 8,407 aSNPs for testing (Denisova n = 2,871; Neanderthal PNG n = 2,867; Neanderthal 1KG n = 2,669), predicted to impact 651 genes. For the ATAC-seq data, this resulted in 1,520 testable aSNPs. We then calculated a moderated version of the log_2_ fold change between introgression carriers and non-carriers using Hedges’ g (since introgressed/non-introgressed state is not consistent across the genome, we cannot use empirical Bayes differential expression frameworks such as Limma-voom; see Methods), and compared this estimate to AlphaGenome predictions of fold change differences between the two alleles. We find that overall, observed Hedges’ g is only weakly correlated with AlphaGenome predictions of RNA-seq (overall Spearman’s *ρ* = 0.012*, p* = 0.270, Figure 1A), whereas prediction of ATAC-seq was more robust (overall Spearman’s *ρ* = 0.111*, p* = 1.1 *×* 10*^−^*^5^, Figure 1B). Considering the three sources of introgressed aSNPs independently, we find that RNA-seq predictions are consistently poor regardless of ancestry source, with no significant correlation between observed and predicted values. ATAC-seq predictions are significant for all three introgression sources after multiple testing correction, suggesting that the ATAC-seq predictions can discriminate instances of differentially active introgression. Although AlphaGenome was trained on GTEx and ENCODE data [22], which are dominated by samples from individuals of European-like genetic ancestries [48], we find no evidence that AlphaGenome predictions are more accurate for Neanderthal 1KG aSNPs than the other two ancestry sources.

**Figure 1.**
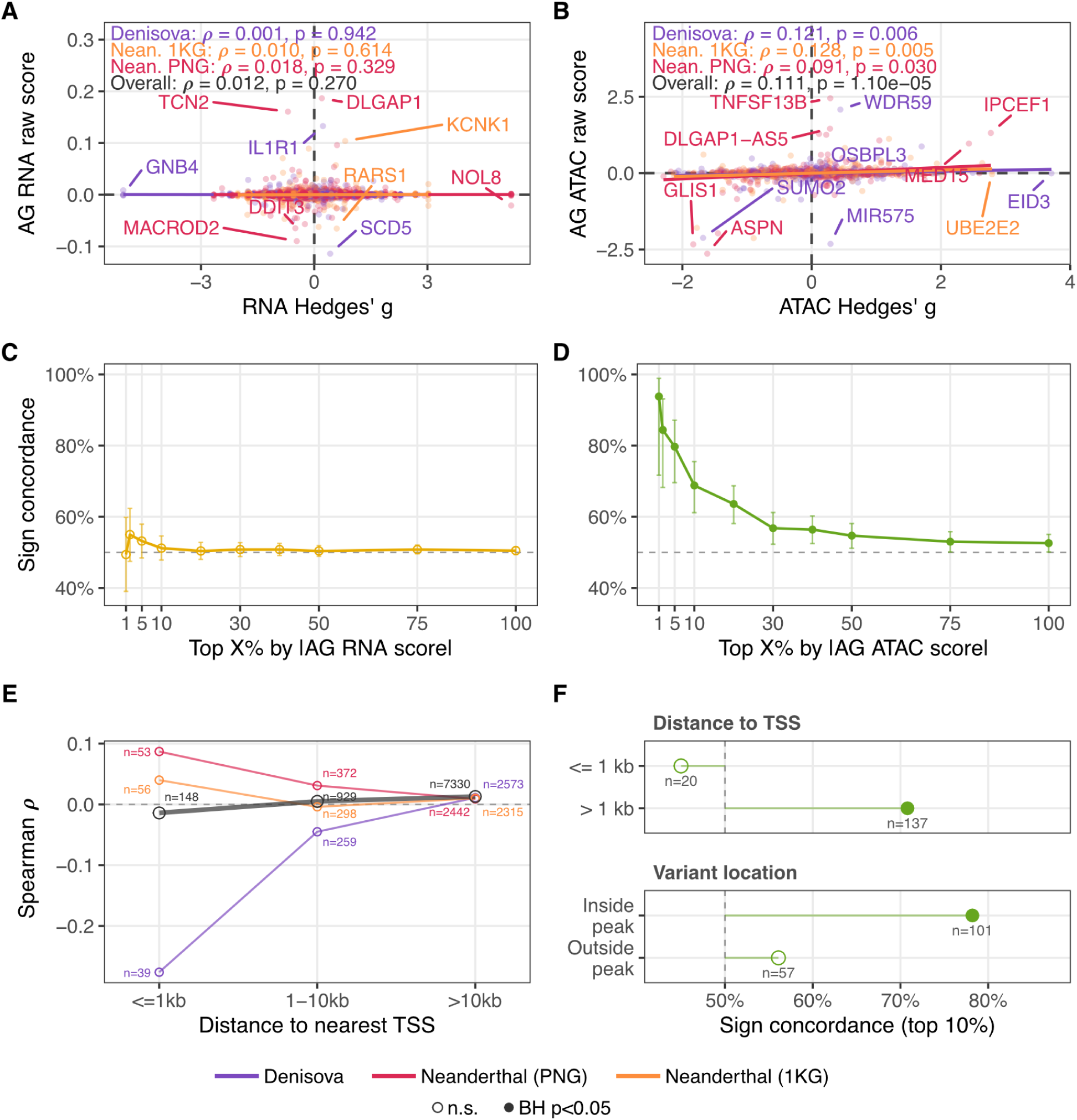
AlphaGenome predicts introgressed allelic effects on chromatin accessibility, but not on expression. **A.** Predicted AlphaGenome RNA-seq log_2_ fold change against observed introgression effect size (Hedges’ g) in PNG LCLs, coloured by introgression source. **B.** As A, for ATAC-seq. **C.** Sign concordance between the AlphaGenome RNA-seq log_2_ fold change and observed effect size in PNG LCLs as a function of predicted effect size magnitude. Error bars are binomial 95% CIs; filled points are significant after BH correction; the dashed line is chance (50%). **D.** As C, for ATAC-seq. **E.** Sign concordance between the AlphaGenome RNA-seq log_2_ fold change and observed effect size in PNG LCLs, stratified by distance to the nearest TSS and by ancestry source. The heavier grey line is pooled across all three sources. **F.** ATAC-seq sign concordance within the top 10% of variants by *|*raw score*|*, stratified two ways: peak annotation (Promoter-TSS vs. other), and whether the variant itself falls inside the peak or only within 250 bp of it. *n* is the number of variants in each top-10% slice. Filled points are significant after BH correction.

The poor concordance between observed and predicted effects on gene expression recapitulate known limitations of S2F models, which struggle to predict allele-specific effects on gene expression levels [27, 22]. We therefore performed a less stringent test, and asked how often the direction of effect agreed between the observed and predicted data. Across the entire dataset concordance between predicted and observed direction of effect for gene expression was no better than random (proportion concordant = 0.505, one-sided binomial *p* = 0.186); this was not the case for chromatin accessibility predictions (overall proportion concordant = 0.526, one-sided binomial *p* = 0.022). We reasoned that this could potentially be due to noise in both the experimental and predicted datasets when effect sizes are small and statistically undistinguishable from zero. Therefore, we stratified data by the absolute magnitude of the Alpha Genome prediction. RNA-seq direction of effect estimates were consistently inaccurate, with no bin performing better than random chance (Figure 1C). In the ATAC-seq data sign concordance between observed and predicted data was higher at the extremes of the predicted AlphaGenome score distribution, and remained higher than random chance across all bins we considered, even as the actual magnitude of the difference between the two alleles in the empirical and predicted data approach 0 (Figure 1D). Stratifying aSNPs by distance to the nearest TSS did not improve the accuracy of sign predictions, regardless of introgression source (Figure 1E), with the correlation between predicted and observed data being worse for aSNPs located within 1 kb of an annotated TSS (although this is dependent on a small number of variants; n = 148). Sign concordance for ATAC-seq predictions was greater when aSNPs were located away from TSS, and when aSNPs were located directly inside an ATAC-seq peak, rather than adjacent to it (Figure 1F).

Although the analyses above used all aSNPs that passed filtering criteria, multiple aSNPs in the same introgressed haplotype frequently have the same nearest TSS. Therefore, we repeated the RNA-seq analyses considering only the largest predicted effect size per gene per ancestry source, for a total of 709 tested variants (a gene can be included more than once if it is the nearest gene to variants across different introgression sources). This has no significant impact in our conclusions (**Supplementary table 4** and **Supplementary table 5**).

### Measures of individual SNP activity match AlphaGenome’s predictions more closely than measures from live cells

Gene expression is the product of multiple cis and trans-acting effects. However, AlphaGenome predictions are limited to assessing the effect of a single SNP at a time. Especially in the case of gene expression, if multiple nearby aSNPs impact expression or chromatin accessibility in an additive manner, examining the AlphaGenome predictions for each constituent aSNP independently will lead to an apparent failure in predictive accuracy. Therefore, we compared activity predictions to the results of a recent massively parallel reporter assay (MPRA) that independently tested both alleles of 17,078 introgressed Denisovan and Neanderthal aSNPs segregating at allele frequencies *>* 0.15 in Papuan individuals [46] for their ability to drive reporter gene expression in an episomal construct. Of these, 1,951 were deemed active, and 180 exhibited differential activity between the introgressed and non-introgressed allele, with no evidence from the MPRA that the remaining 15,126 meaningfully drove gene expression *in vitro*. The MPRA readout is expression of a reporter gene construct, which AlphaGenome cannot directly predict; therefore we considered both RNA and ATAC-seq fold change predictions in the analyses below.

As above, we found that AlphaGenome predictions for RNA-seq were more discordant than the ATAC-seq predictions (Spearman *ρ* = 0.072*, p* = 1.7 *×* 10*^−^*^3^ for RNA-seq; Spearman *ρ* = 0.230*, p* = 9.3 *×* 10*^−^*^25^ for ATAC-seq, Figure 2A), with results again being comparable across introgression sources. Correlation between observed and predicted scores improved when we considered only aSNPs classified as differentially active in the MPRA, where experimental log_2_ FC estimates are larger (Spearman *ρ* = 0.155*, p* = 3.8 *×* 10*^−^*^2^ for RNA-seq; Spearman *ρ* = 0.435*, p* = 1.0 *×* 10*^−^*^9^ for ATAC-seq, Figure 2A), although the confidence intervals for RNA-seq estimates spanned 0. Sign concordance between predicted and observed values for differentially active variants was significant for ATAC-seq (proportion concordant = 0.672, one-sided binomial *p* = 2.2 *×* 10*^−^*^6^) and RNA-seq (proportion concordant = 0.567, one-sided binomial *p* = 0.042), and concordance was higher at aSNPs where AlphaGenome predicted a greater effect size, confirming the hypothesis above (Supplementary Figure 2). Furthermore, when we stratified the MPRA aSNPs by activity level (that is, inactive, active, and differentially active) we found that AlphaGenome scores were systematically different between the three classes across both data modalities (Supplementary Figure 3).

**Figure 2.**
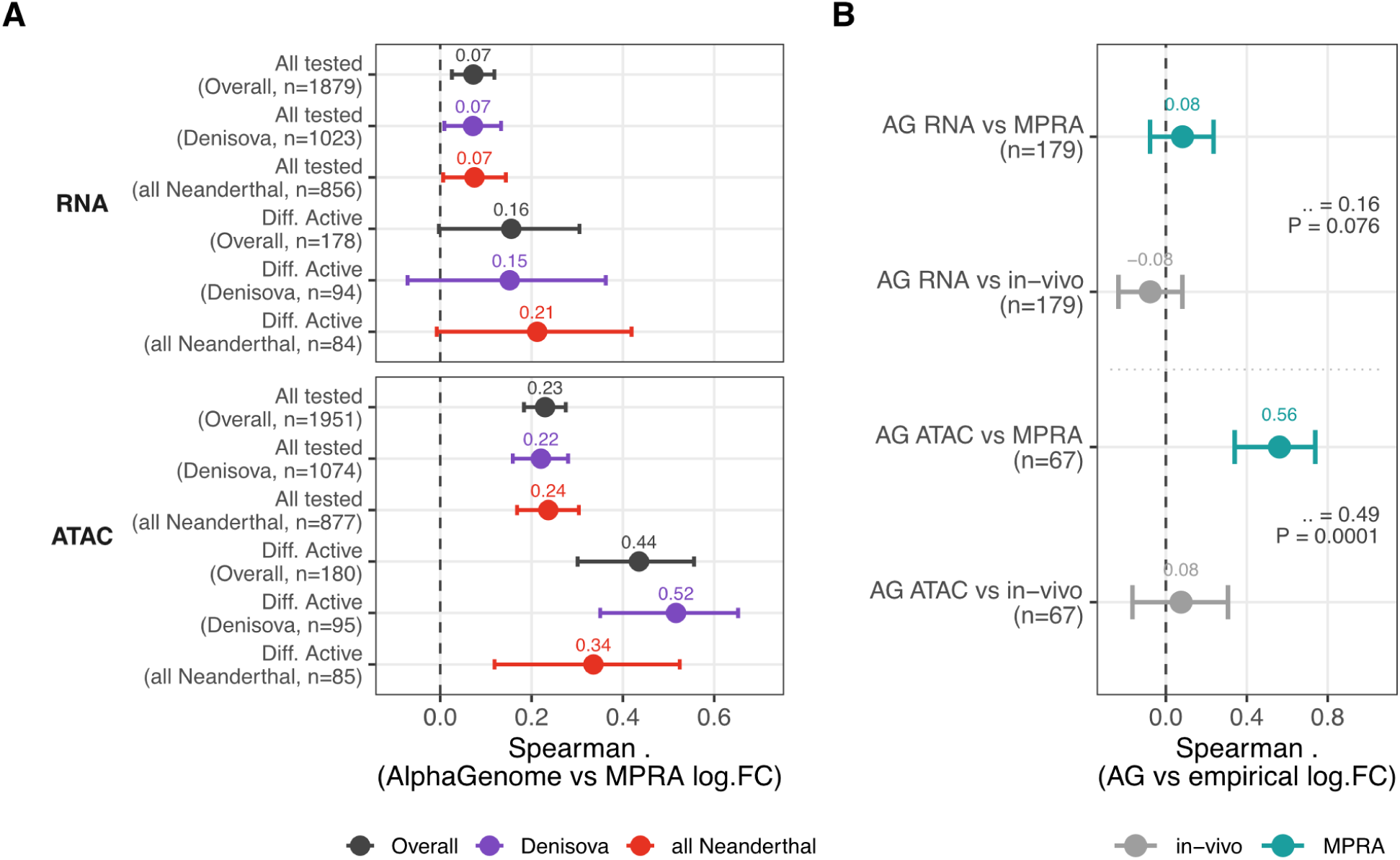
AlphaGenome predictions track MPRA-measured allelic activity more closely than they track effects measured in live cells. **A.** Absolute AlphaGenome quantile score in GM12878 for MPRA-tested introgressed variants, grouped by their measured MPRA activity class and split by introgressed ancestry source. **B.** Spearman *ρ* between the AlphaGenome raw score and each of two empirical log_2_ fold-change estimates for the same variants: in-vivo estimate from LCL data (grey) and the MPRA estimate from Comerford et al. (teal). Error bars are bootstrap 95% confidence intervals from 10,000 resamples.

A small number of variants were tested in the MPRA and also part of the datasets that underpin Figure 1 (RNA-seq n = 179, ATAC-seq n = 67). For these we asked which type of experimental readout better correlated with AlphaGenome scores (Figure 2B). Correlation of either experimental measurement with RNA-seq predictions was no better than chance, but predicted ATAC-seq measurements correlated far better with the MPRA allelic fold change estimates (Spearman *ρ* = 0.562) than with peak measurements in experimental ATAC-seq data (Spearman *ρ* = 0.075), again highlighting that AlphaGenome’s predictions recapitulate the effects of individual aSNPs. The small number of aSNPs included in these comparisons was partly due to the allele frequency threshold we applied to the data in Figure 1, which requires that at least three individuals carry at least one copy of the introgressed allele. Since RNA-seq data was available only from 9 individuals, this was equivalent to an introgressed allele frequency threshold *≥* 0.167, slightly higher than the 0.15 cut-off from [46]. Relaxing this filter to consider all sites where at least two individuals carried at least one copy of the introgressed allele (equivalent to an introgressed allele frequency cut-off *≥* 0.111) resulted in a significant increase in testable sites, but did not impact overall trends (Supplementary Figure 4).

### Extending predictions to multiple tissue types

The analyses above demonstrate that AlphaGenome’s chromatin accessibility predictions correlate with experimental observations. However, experimental data is limited to that generated from LCL lines, while introgression has been shown to impact multiple cell types, both immune and non-immune. Thus we generated AlphaGenome ATAC-seq and RNA-seq predictions across all 144,139 aSNPs for an additional set of 14 tissues, including two additional immortalised cell lines, as well as five additional tissues where only RNA-seq was available, to examine trends across multiple tissues (**Supplementary table 1**; AlphaGenome cannot currently predict ATAC-seq tracks for any neural tissue). These additional Biosamples were selected to provide a broad representation of cell types and tissues previously described as likely harbouring varying proportions of functional introgression [13]. In order to directly compare between Biosamples and data modalities we followed recommendations in AlphaGenome’s documentation and consider only reported quantile scores, *q*, in the following sections (see Methods and [22]).

Principal component analysis of AlphaGenome predictions demonstrated that these carry a degree of tissue specificity. Considering both RNA-seq and ATAC-seq predictions for either differential activity or overall activity levels we observed generally similar trends (Figure 3A and Supplementary Figure 5), with the first two principal components separating immune and non-immune samples, as well as primary samples from the three immortalised cell lines. In the case of RNA, the second principal component additionally separated the neural samples. Once again, these effects were consistent across the three archaic ancestries (Supplementary Figure 6), suggesting that aggregation across Biosamples and introgressed regions both primarily recapitulates known principles of chromatin organisation genome-wide [49]. However, given the poor correlation between the experimental and predicted RNA-seq measurements (figures 1 and 2), we elected to carry out the analyses below only on the ATAC-seq log_2_ fold change predictions.

**Figure 3.**
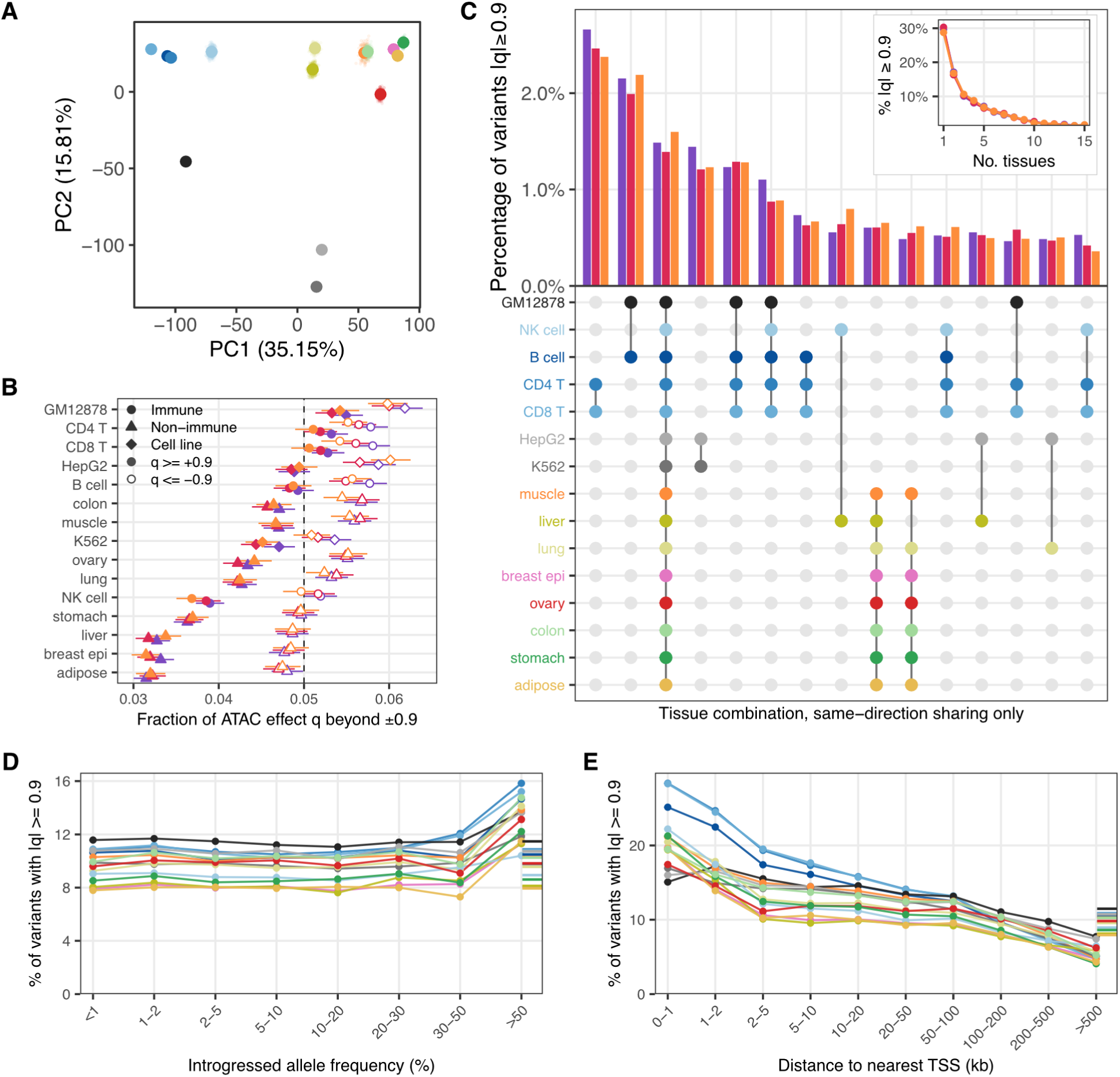
AlphaGenome predicts a broadly cross-tissue regulatory landscape for introgressed variants. **A.** Principal component analysis of AlphaGenome ATAC-seq effect scores across Biosamples, from 500 haplotype-thinned draws Procrustes-aligned behind their mean. Colours denote distinct Biosamples as labelled in panel C. **B.** Fraction of introgressed variants with *|q| ≥* 0.9 in each Biosample, coloured by introgression source and stratified by direction of effect. Error bars are exact binomial 95% CIs. **C.** Top 15 most frequent tissue combinations among strong calls, as a fraction of each source’s strong set; dots mark the member tissues, ordered by clustering on the between-tissue effect correlation. Inset: number of tissues per strong variant. **D.** Percentage of introgressed variants reaching *|q| ≥* 0.9 in each Biosample, binned by introgressed allele frequency. One line per Biosample; the rug on the right marks each Biosample’s own overall rate. Colours denote distinct Biosamples as labelled in panel C. **E.** As D, binned by distance from the variant to its nearest TSS.

First, we examined the absolute distribution of quantile scores. 50,010 aSNPs had a *|q| ≥* 0.9 in at least one Biosample. Considering each Biosample separately, seven had a significant excess of extreme quantile aSNPs relative to expectations, and this excess ranged from 1.15x in GM12878 to 1.03x in colon. However, more introgressed alleles had a negative score (*q ≤ −*0.9) than a positive one (*q ≥* 0.9, Figure 3B); that is, AlphaGenome consistently predicted that introgressed alleles at the extremes of the quantile score distribution decrease chromatin accessibility more frequently than they increase it. Across the 15 biosamples only GM12878, CD4^+^ and CD8^+^ T cells—all immune Biosamples—had a significant excess of aSNPs where the introgressed allele was predicted to increase accessibility; while significant in some Biosamples, the magnitude of this excess amounted to no more than a 1.08x fold increase. Only adipose tissue, breast epithelium and liver predictions were depleted of both extremely positive and negative effects. These overall trends of stronger excesses in either direction in immune or immune-like Biosamples, and the depletion of extreme quantile variants in tissues such as liver or breast epithelium, were generally robust to varying the quantile score threshold (Supplementary Figure 7). When considering introgression sources, the excess (in either direction) was most pronounced in Denisovan aSNPs, followed by Neanderthal PNG aSNPs and finally in Neanderthal 1KG aSNPs, although the magnitude of these differences was consistently small, and did not reach significance in any Biosample (all BH-adjusted *p >* 0.12).

We then examined patterns of signal sharing across tissues in greater detail. 14,818 aSNPs (29.6% of aSNPs with an extreme quantile score in at least one Biosample) were predicted to impact only a single Biosample; these were concentrated in the three immortalised cell lines and skeletal muscle, and least common in adipose tissue (**Supplementary table 6**). Sharing of extreme signals with concordant direction of effect across Biosamples was common, and patterns of sharing primarily reflect how densely our predictions sampled different developmental lineages (e.g. immune and immune-like Biosamples, HepG2 and liver; Figure 3C); again confirming that AlphaGenome’s predictions can recapitulate true regulatory structure. Only 742 aSNPs with a score *≥* 0.9 had the same sign in all 15 Biosamples; although this number is small, it was the third most common multi-Biosample configuration in the dataset.

*|q| ≥* 0.9 aSNPs were not randomly distributed across introgressed haplotypes with regard to either allele frequency or distance to a TSS. We observed an excess of introgressed alleles with frequencies *≥* 0.5 and extreme quantile scores across all Biosamples (Figure 3D). This excess was significant in 12 Biosamples (BH-corrected Fisher’s exact *p <* 0.05 in 12 out of 15), ranging from 1.48x in lung to 1.27x in HepG2, and did not reach significance only in K562, GM12878 and NK cells. Only 1,105 aSNPs have an introgressed AF *>* 0.5; of these, 463 reach *|q| ≥* 0.9 in at least one Biosample, and they sit at a median distance of 53.4 kb from their nearest TSS, with only 81 within 10 kb. We also observed structure when considering the relationship between *|q|* and distance to the nearest TSS (Figure 3E). aSNPs located closer to a TSS tended to have higher quantile scores, likely reflecting the regulatory architecture of the genome, where promoters are located closer to a transcription start size and tend to have larger effects on gene regulation than distal enhancers [50]. Again, the absolute number of TSS-proximal aSNPs is small (1,612 aSNPs are located within 1 kb of a TSS, and 12,917 within 10kb), but the excess is significant across Biosamples (BH-corrected Fisher’s exact *p <* 0.05 in all 15), ranging from 2.62x in CD8^+^ T cells to 1.31 in GM12878. Three immune cell types (CD8^+^ T, CD4^+^ T, and NK cells) have the largest excess, and the three immortalised cell lines the smallest.

### Prioritising candidates for functional introgression with AlphaGenome

Having shown that there is tissue specificity in the ATAC-seq predictions, we then asked whether we could use this signal to directly predict the functional consequences of introgression. With a median length of 55.2 kb, introgressed haplotypes span between 1 and 26 genes. Distinguishing which, if any of these, is impacted by introgression remains challenging in the absence of direct experimental validation. A possible solution would be to identify which, if any, variants within a haplotype was most likely to have functional consequences. We reasoned that the quantile score served as a good proxy for this. However, as shown above, extreme quantile aSNPs are not independently distributed across the genome nor across haplotypes, with an excess of signal near transcription start sites, and at aSNPs with higher introgressed allele frequencies.

Ideally, we would have used the RNA-seq predictions to directly identify the specific gene or genes impacted by a given aSNP, by examining the predicted magnitude of the log_2_ fold change between the introgressed and non-introgressed alleles at each gene with a score prediction. However, raw RNA-seq AlphaGenome log_2_ fold change predictions are very compressed: 98.3% of predictions had a absolute log_2_ fold change *<* 0.01, and the broadest IQR amongst the twenty Biosamples with RNA-seq predictions was 0.0020, in gastrocnemius medialis. When transformed into quantile scores these small differences become artificially magnified, such that ATAC-seq and RNA-seq quantile scores for introgressed aSNPs were not well correlated (Spearman *ρ* across all Biosamples together = 0.075). Nevertheless, we observed a clear excess of variants with extreme quantile scores in both the ATAC-seq and RNA-seq predictions (Supplementary Figure 10), suggesting that true signal may reside at the extremes of either distribution. This observation was supported by the fact that, as shown in Figure 1 and Supplementary Figure 2, concordance between empirical and predicted direction of effect was higher at more extreme values of *q*.

One possibility is that introgression that survives to this day preferentially impacts certain genes or pathways. Therefore, we performed gene enrichment testing for each Biosample across 20 *|q|* and distance to nearest TSS thresholds (Figure 4A; see Methods), a total of 240 enrichment tests. For each test, we compared genes where at least one aSNP was located within the distance threshold and exceeded the *|q|* threshold, relative to all genes located within the distance threshold. By sweeping across *|q|* and distance thresholds we were able to identify terms that consistently reoccurred, rather than those that appeared only once or twice, and examine how the choice of threshold impacts the results. A total of 140 terms (9 Hallmark gene sets, and 131 GO terms) were significantly overrepresented across the range of thresholds and Biosamples, 57 of them more than once. Of these, the Hallmark term “TNFA signaling via NFKB” was the most consistent signal, reaching significance in 32 different tissue*×|q|×*distance combinations after multiple testing correction (**Supplementary table 7**). Enrichment in this pathway occurred primarily in the immune Biosamples, and was driven by 47 distinct genes across a range of distances and *|q|* thresholds (Figure 4B). The TNFA pathway modulates the activation of the adaptive immune system through NF-*κ*B signalling, and individual members, such as *TNFAIP3* have been previously shown to harbour introgressed variants that impact immune system function in present-day individuals [51]. Multiple additional immune pathways, such as interferon gamma and interleukin signalling pathways were also enriched, although less consistently than TNFA signalling. Other pathways that reoccurred often include Hallmark “UV response (down)”, which was significant in 8 different tissues and 14 different combinations, and a set of nested GO terms associated with protein phosphorylation, significant in 17-19 Biosample*×|q|×*distance thresholds. We then tested if the enrichments could be reproduced by testing only one introgression source at a time, we found that no enrichment remained significant. Gene foreground and background sets for each ancestry are small, however, such that we do not have sufficient statistical power to determine whether this is a true negative result, or due to only a small number of testable genes.

**Figure 4.**
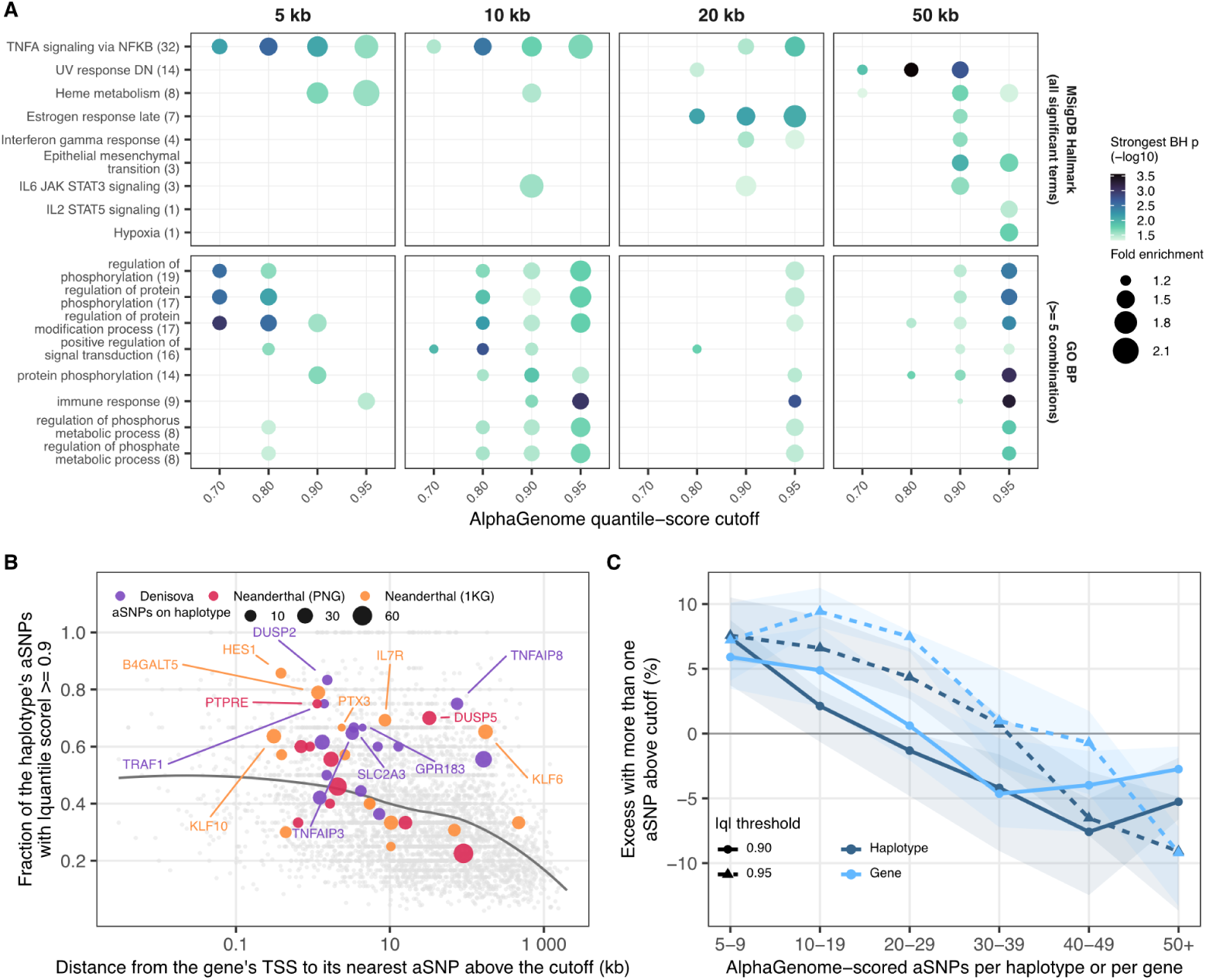
Extreme AlphaGenome quantile scores cluster within introgressed haplotypes. **A.** Gene-set over-representation analysis across a grid of distance-to-TSS and *|q|* thresholds. All nine significant Hallmark terms, and GO BP terms reaching BH *<* 0.05 in at least five of the 20 threshold combinations are shown. The parenthetical after each term indicates the number of significant Biosample *×* threshold combination tests it was significant in, of 240 possible. The 1 kb window returned no significant terms at any cut-off and is not shown. **B.** Median haplotype distance to the TSS against the fraction of haplotype aSNPs with *|q| ≥* 0.9 for the 39 TNFA/NF*κ*B genes significant in at least four tests. **C.** Observed minus expected probability that a haplotype or gene carries more than one aSNP above a *|q|* cut-off, given that it carries at least one, against the number of AlphaGenome-scored aSNPs it contains. The expectation is the Poisson-binomial over each aSNP’s own predicted probability, ribbons are the 10th–90th percentile across the 12 primary tissue Biosamples.

As an orthogonal approach, we considered each haplotype with at least one *|q| ≥* 0.9 aSNP and asked whether it had more such aSNPs than we would predict by chance after controlling for the number of SNPs in the haplotype (Figure 4C). Using two different approaches, one of which additionally controlled for the overall excess of signal near TSS (see Methods), we consistently observed that haplotypes containing up to 20 aSNPs were significantly enriched for runs of extreme *|q|* aSNPs. We repeated these analyses setting the *|q|* threshold to 0.7, 0.8 or 0.95 (Supplementary Figure 11), and found that at the more stringent cut-off, the excess remained significant for haplotypes containing up to 39 aSNPs, but decayed as the number of SNPs in a haplotype increased beyond that number. This breakdown in signal likely reflects genuine signal driven by the local regulatory context: as haplotypes get longer they are less likely to share a single regulatory state. Together, these results suggested that while pinpointing a singular functional variant within a haplotype is likely not currently possible with AlphaGenome scores, there might be information in the fraction of aSNPs within a haplotype that exceed a given *|q|* threshold, and that AlphaGenome predictions may provide some degree of functional information as to where in a haplotype functional variants are most likely to reside, even though its predictions are strictly made at the individual aSNP level.

Therefore, we examined whether these patterns of excess varied across Biosamples. Of 60,341 testable Biosample*×*haplotype combinations at *|q| ≥* 0.9, our Poisson-binomial approach detected an excess of extreme *|q|* scores in 186 different haplotypes (369 Biosample*×*haplotype combinations, Supplementary Figure 12, **Supplementary table 3** and **Supplementary table 8**). When we consider the 47 haplotypes with a significant excess of extreme *|q|* scores in at least three Biosamples (Figure 5), we find that these can be clustered into three distinct categories—haplotypes where the signal is predominantly restricted to immune Biosamples, including genes such as *JAK1* or *TAB2*, haplotypes where the signal is broadly shared across all tissues, and a small set of haplotypes where the signal clusters across all non-immune tissues; Supplementary Figure 13 shows examples of haplotypes driven primarily by immune and non-immune patterns. This was the only analysis where we observed significant differences between the three ancestry sources: Neanderthal 1KG haplotypes were less likely than Neanderthal PNG or Denisovan haplotypes to be associated with extreme scores (BH-corrected Fisher’s exact *p* = 0.036). Surprisingly, we observed no association between introgressed allele frequency and how often haplotypes were deemed significant in this approach (Spearman *ρ* = 0.014*, p* = 0.21). We used similar logic to detect regions with a deficit of extreme *|q|* scores, and identified 16 significant regions at *|q| ≥* 0.9 (Supplementary Figure 12, **Supplementary table 2** and **Supplementary table 8**). These regions were biased towards longer haplotypes with a higher number of SNPs due to the difficulty in identifying a deficit of extreme signals in small haplotypes **Supplementary table 2**, but they nonetheless suggest that a combinatorial approach examining both the excess and depletion of strongly scoring variants in archaic haplotypes may provide a more complete picture of the functional archaic introgression landscape.

**Figure 5.**
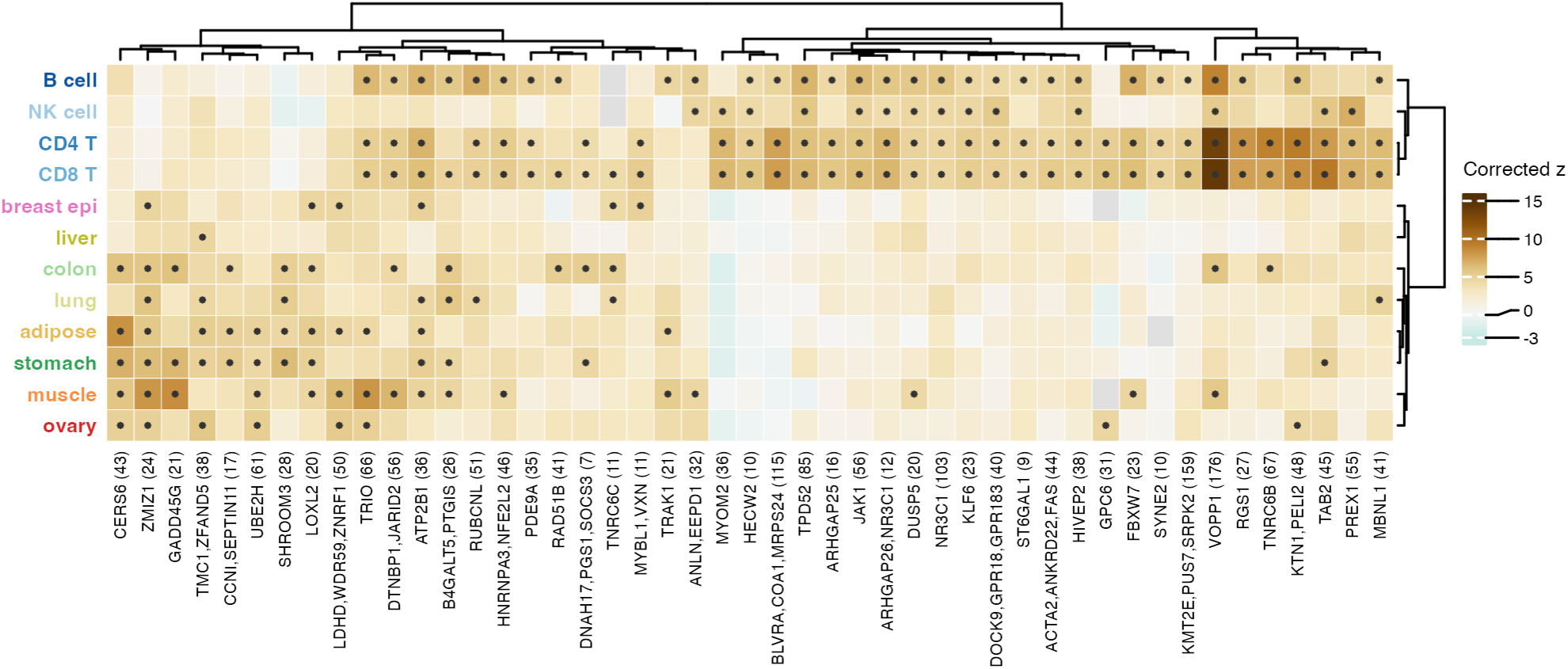
Haplotypes with a recurrent excess of extreme predictions separate into immune and non-immune sets. Standardised excess of aSNPs with *|q| ≥* 0.9 per haplotype and Biosample, for the 47 haplotypes reaching BH *<* 0.05 in at least three of the 12 primary Biosamples. Points mark BH-corrected *p <* 0.05 in that Biosample; grey cells are haplotypes with no aSNP above the cut-off in that Biosample. Biosamples clustered on pairwise-complete Spearman correlation. Numbers inside the parenthesis are the number of aSNPs within the haplotype

## Discussion

Understanding the functional consequences of archaic hominin introgression remains challenging. Sequence-to-function models can fill this gap, but their full utility for interrogating archaic hominin introgression remains unclear. Here, we have assessed the accuracy and potential of AlphaGenome for understanding introgressed variant effect by generating predictions for over 144,000 introgressed variants and comparing these predictions to three modalities of experimental functional genomics data – RNA-seq, ATAC-seq and an episomal MPRA readout. We focused on individuals of Papuan-like genetic ancestry because while they have some of the highest levels of archaic introgression worldwide [52], functional genomics data from these populations remains extremely limited both in terms of the number of samples analysed and tissue breadth. This rules out using publicly available datasets to characterise introgressed variants segregating in these individuals.

AlphaGenome’s predictions frequently corroborated findings made using orthogonal approaches. For instance, multiple groups have documented signals of adaptive introgression near immune genes, especially in present-day Papuan individuals [53, 54, 13, 32]. Here we show that these introgressed blocks are predicted by AlphaGenome to impact immune gene expression rather than the expression of genes in other tissues. Our results suggest that introgression appears to have specifically targeted multiple genes in the TNF signalling pathway, and, to a lesser degree, in other adaptive immune system signalling pathways. However, contrary to previous reports, we did not observe any marked differences in predicted effects across introgression sources (Neanderthals vs Denisovans). It is unclear if this is due to poor resolution in the AlphaGenome predictions, or if this reflects the fact that previous reports incorporated less nuanced predictions of functional impact. Likewise, it is not immediately obvious why AlphaGenome’s predictions of ATAC-seq data should better recapitulate observed MPRA results than its RNA-seq predictions. A possible explanation is that although the readout of an MPRA is expression of a reporter construct, episomal MPRAs assess the regulatory potential of tested variants, and ATAC-seq data may be a better, or more direct proxy for that measurement.

Although AlphaGenome predictions were ultimately sufficiently accurate, especially when including ATAC-seq data, to infer signal at extreme quantile score values, the challenge of linking individual aSNPs to the gene they impact remains unsolved. Using the simple heuristic of linking aSNPs to their nearest transcription start site (TSS), we found significant enrichment in immune pathways. More sophisticated methods that directly make use of experimental data, such as E2G [55], would likely help refine this assignment and clarify these signals, but would not address the general weakness of RNA prediction quality across models, or the compressed range that characterised the RNA-seq raw score predictions. Further, additive prediction of haplotype effects, as done by prediXcan and JTI [56, 57], is not directly possible with AlphaGenome at present, again limiting linkages between predicted effects and phenotypic outcomes. Similarly, AlphaGenome predictions reflect a Biosample in its steady state, which especially in the case of immune cells, may not capture the full regulatory potential of introgressed sequences. Robust prediction of cellular states after perturbation remains a goal for S2F models [58, 59].

RNA-seq predictions, which in general are much more readily ‘understandable’ in terms of phenotypic outcome, were significantly less accurate that predictions of local ATAC-seq levels. This is a well-documented failure of S2F models in general [26, 27], and our results exacerbate the impression of poor performance at this task. The original AlphaGenome publication reported high concordance between predicted and observed direction of effect for fine-mapped eQTLs [22]. This result is markedly different from what we report, but the discrepancy is readily explained: fine-mapped eQTLs have a high prior probability of measurably impacting gene expression at one or more nearby genes, an assumption that is not necessarily met by our set of introgressed SNPs. It is also likely that some of the poor correlations we observe between experimental and predicted data reflects true biology: most introgressed aSNPs are likely to have little to no impact on gene expression levels. Finally, our experimental RNA-seq and ATAC-seq datasets were derived from relatively few individuals, limiting our power to detect small effects, and in turn making it harder to assess the robustness of the predictions of AlphaGenome. In this context, others have shown that the accuracy of ChromBPNet predictions appear to be independent of allele frequency [60], which suggests that S2F models hold great potential for predicting the effect of rare regulatory variants that are hard to quantify using observational data from human carriers. Overall, our results suggest that AlphaGenome can identify meaningful trends and extend current knowledge on the consequences of introgression, but significant caution is required in interpreting its output.

## Acknowledgements

We thank sample donors at the University of Papua New Guinea, and members of the Gallego Romero lab for comments. This work was supported by awards from the Australian Research Council (DP200101552) to IGR; the Marsden Fund of the Royal Society of New Zealand (20-MAU-017) to MPC, IGR, and NB; the National Health and Medical Research Council of Australia (#2042468) to TMJ; the French National Research Agency (ANR) (PAPUAEVOL ANR-20CE12-0003-01) to FXR; the Occitanie region, the European Regional Development Fund (ERDF), and the French government through the France 2030 projects managed by the National Research Agency (ANR-22-EXES-0015 PAPALIM and ANR-24-CE21-0317-01 NUTRIOCEO) to NB; and awards from the Estonian Research Council to MD (TK214) and to MD, DY, and IGR (PRG3113). St Vincent’s Institute acknowledges the infrastructure support it receives from the National Health and Medical Research Council Independent Research Institutes Infrastructure Support Program and from the Victorian Government through its Operational Infrastructure Support Program. The funders had no role in study design, data collection and analysis, decision to publish, or preparation of the manuscript.

## Author contributions

- Conceptualization: MC, IGR
- Formal analysis: MC, IGR
- Funding Acquisition: IGR, FXR, MPC, NB
- Data Curation: MC, IGR
- Investigation: MC, IA, TJ, IGR
- Methodology: MC, IGR, IA, TJ
- Resources: ML, CK, FXR, NB, DY, MD
- Supervision: RA, MPC, IGR
- Visualization: MC, IGR
- Writing – Original Draft Preparation: MC, IGR
- Writing – Review & Editing: All

## Supplementary Figures

**Supplementary Figure 1.**
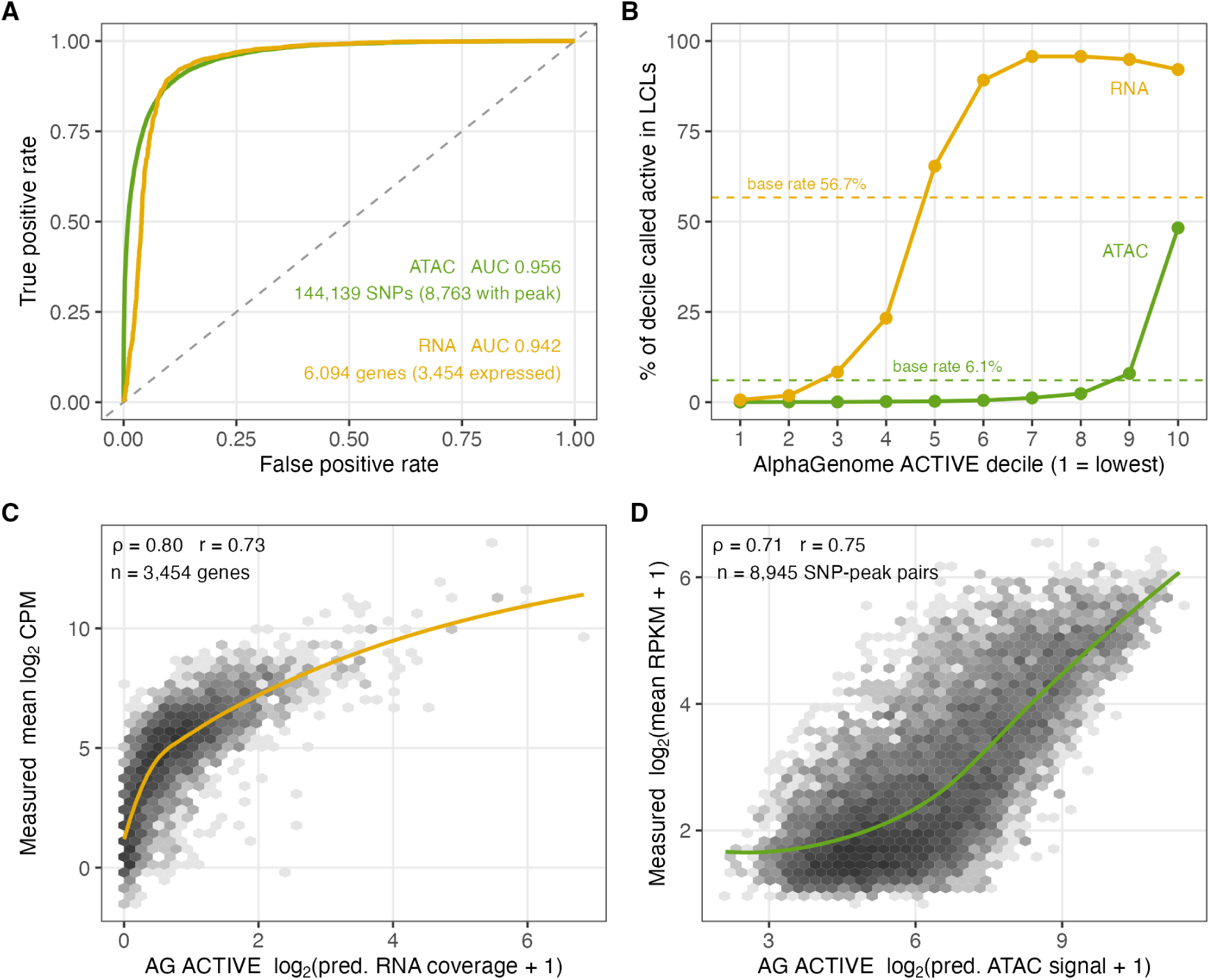
AlphaGenome’s baseline predictions of expression and accessibility are both accurate in lymphoblastoid cells. **A.)** ROC curves for the AlphaGenome ACTIVE scores predicting measured activity in PNG LCLs: for RNA-seq, whether a gene is called expressed; for ATAC-seq, whether a called peak overlaps the variant’s 501 bp scoring window. **B.)** Percentage of observed features (genes or peaks) in PNG LCLs as a function of AlphaGenome’s ACTIVE predictions. The dashed line represents the sample-wide expectation. **C.)** Measured mean expression in PNG LCLs against AlphaGenome ACTIVE RNA-seq score, one point per gene. Trend line is a loess fit. **D.)** As C, for ATAC-seq: measured mean accessibility against the ACTIVE ATAC-seq score, one point per scored windows. Variants whose scoring window spans two adjacent peaks contribute a point for each.

**Supplementary Figure 2.**
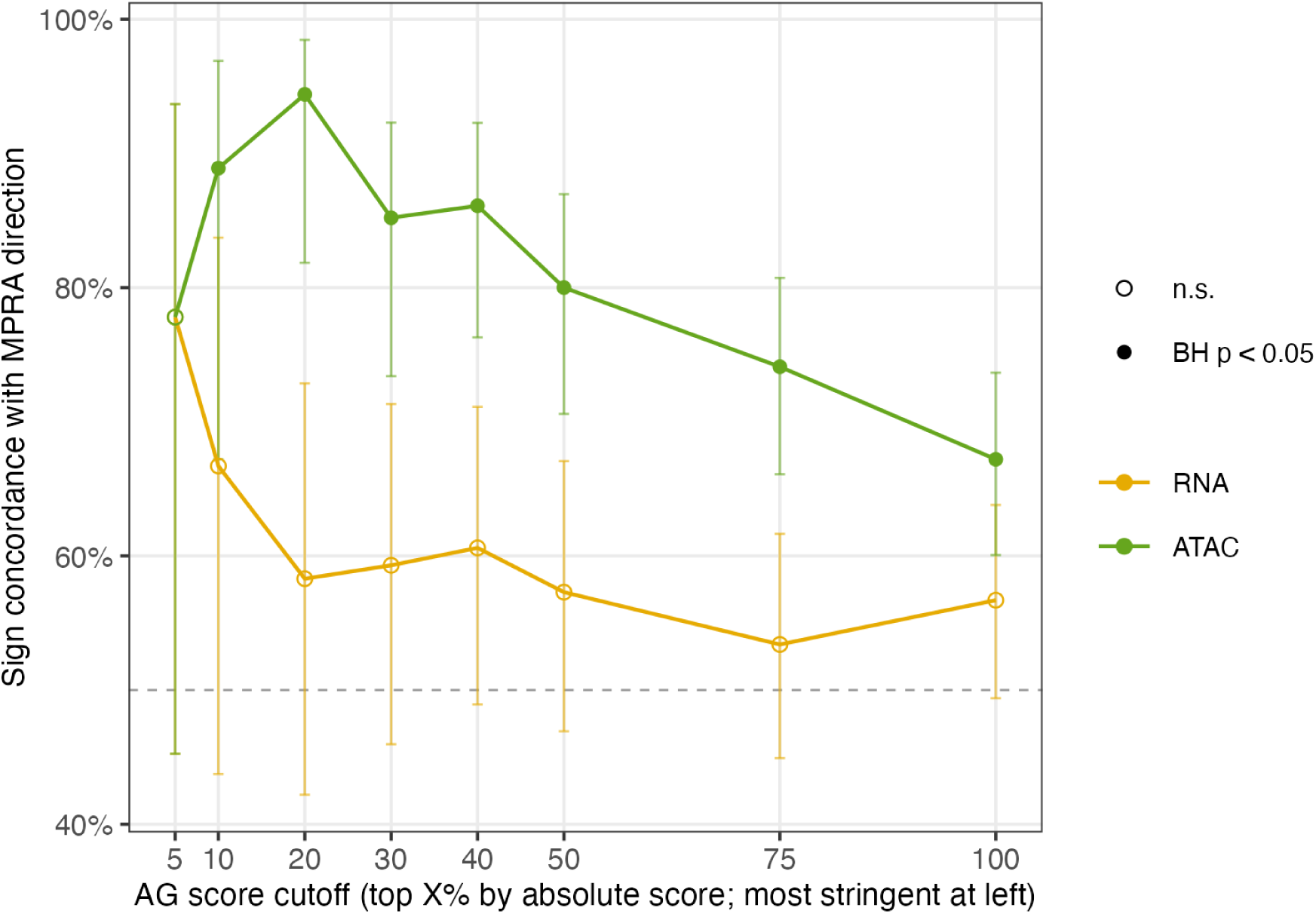
Sign concordance between AlphaGenome predictions and the MPRA-measured direction of effect across all differentially active MPRA variants. Error bars are Wilson 95% CIs, filled points are significant after BH correction across the sweep, and the dashed line is chance (50%).

**Supplementary Figure 3.**
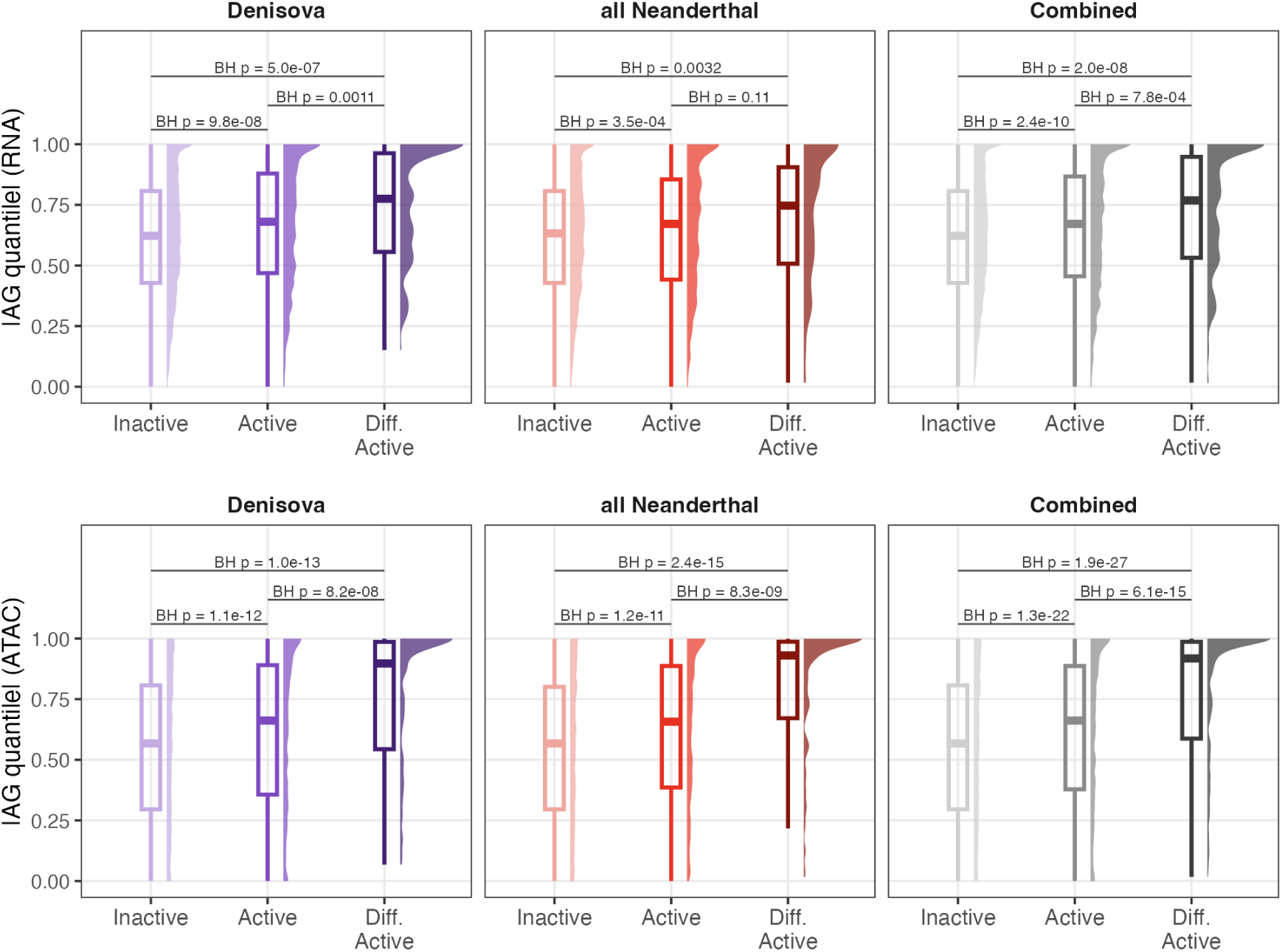
Absolute AlphaGenome quantile score in GM12878 for MPRA-tested introgressed variants, grouped by measured MPRA activity class. Brackets give BH-adjusted pairwise *p*-values.

**Supplementary Figure 4.**
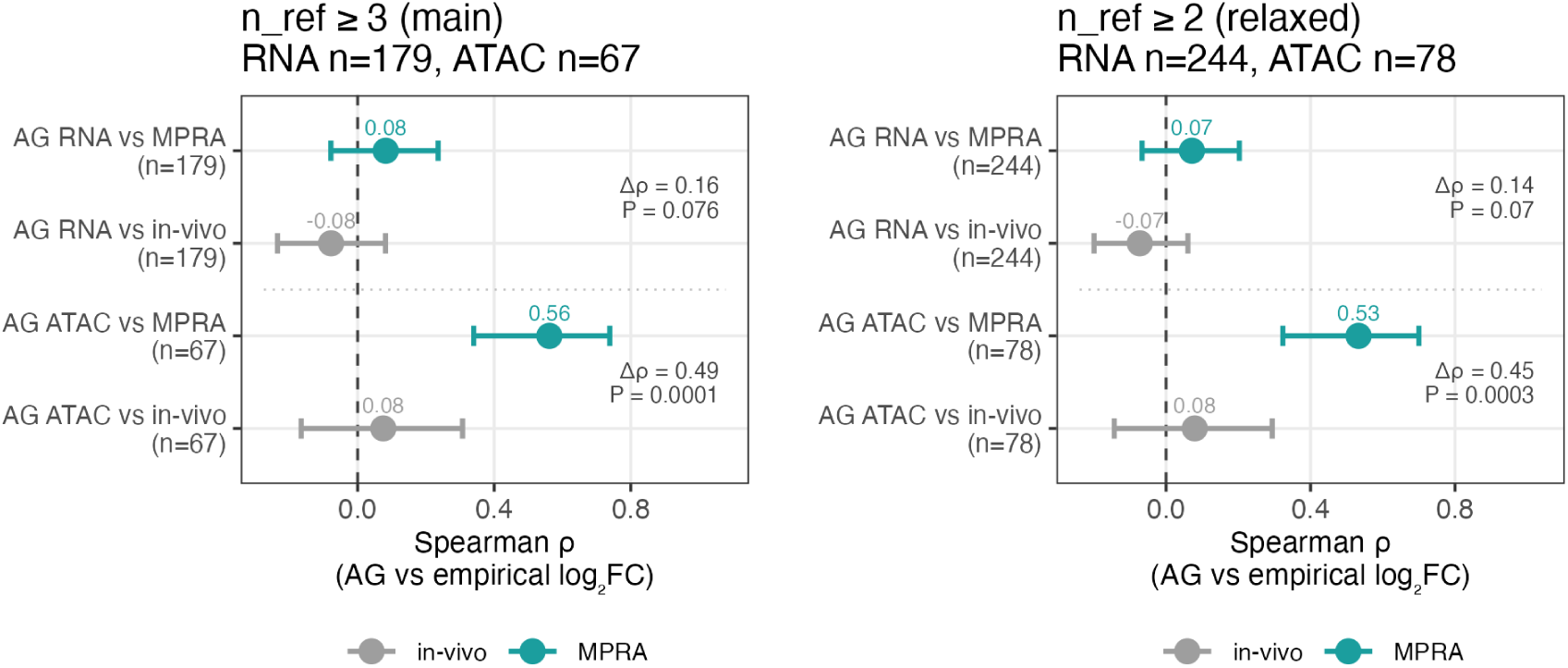
Spearman *ρ* between AlphaGenome effect size raw scores and estimates from LCL data (grey) and the MPRA estimate (teal) at two introgressed allele frequency carrier thresholds, *n* = 3 and *n* = 2. Error bars are bootstrap 95% CIs from 10,000 resamples.

**Supplementary Figure 5.**
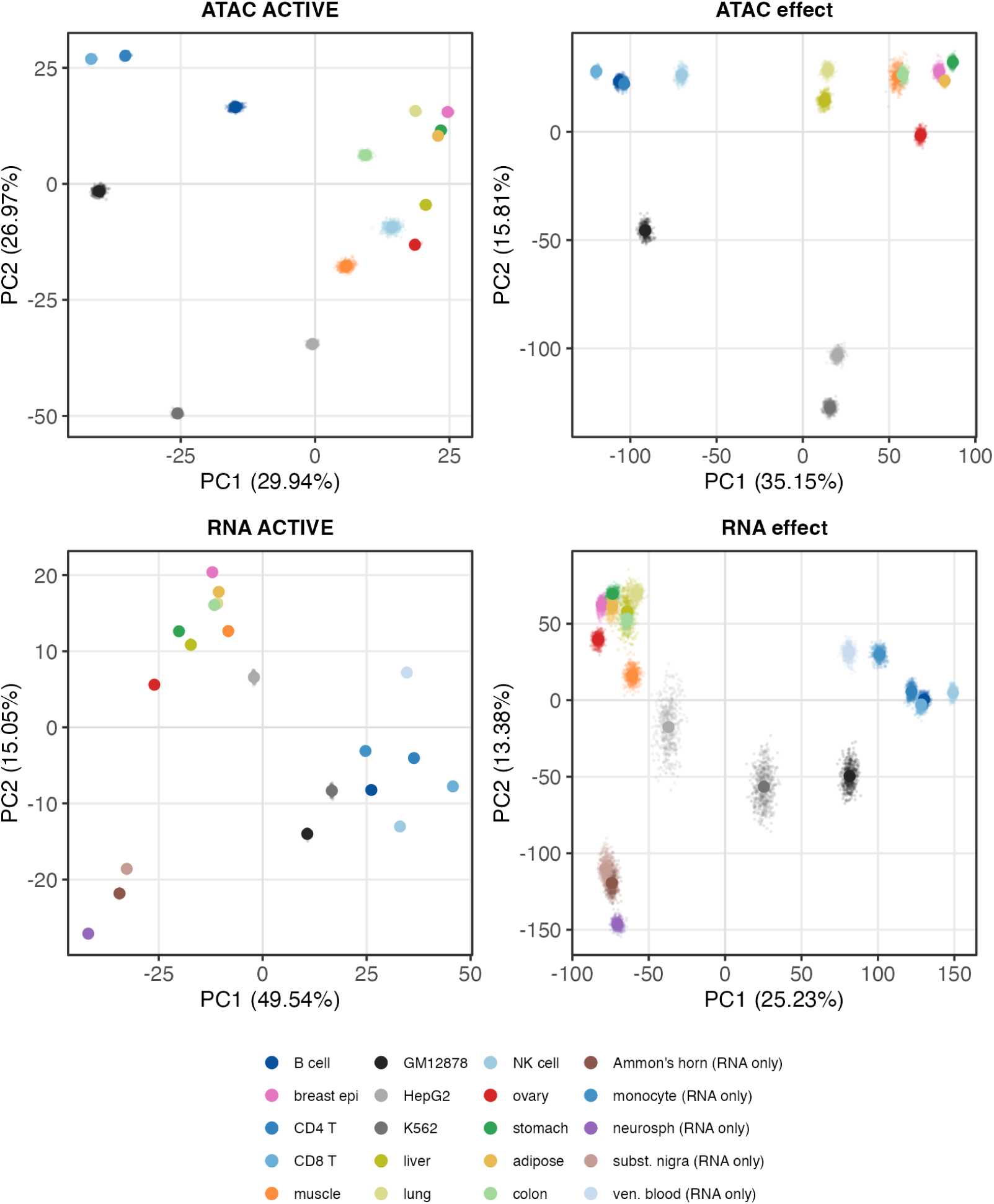
Procrustes-aligned principal component analysis of AlphaGenome predictions across Biosamples, for each of the four score types: ATAC-seq ACTIVE, ATAC-seq effect, RNA-seq ACTIVE and RNA-seq effect. Each panel shows 500 haplotype-thinned draws as faint points behind their mean, one point per Biosample.

**Supplementary Figure 6.**
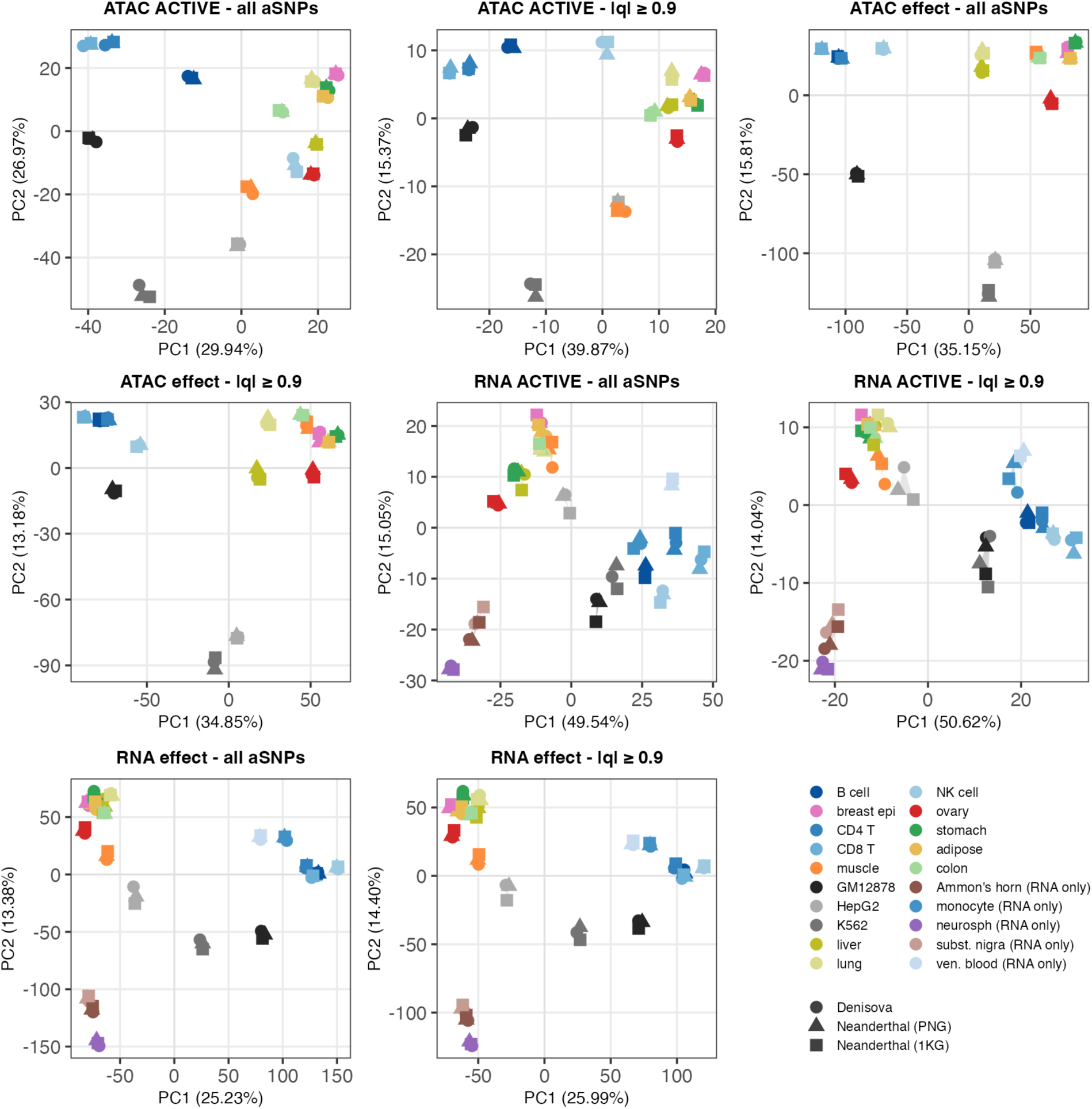
Procrustes-aligned principal component analysis of AlphaGenome predictions across Biosamples, computed separately within each introgression source. Plots on the left column use all scored variants from each score type, plots in the right column use only variants with *|q| ≥* 0.9 in at least one Biosample, defined within each source. Introgressed ancestry source is indicated by different point shapes.

**Supplementary Figure 7.**
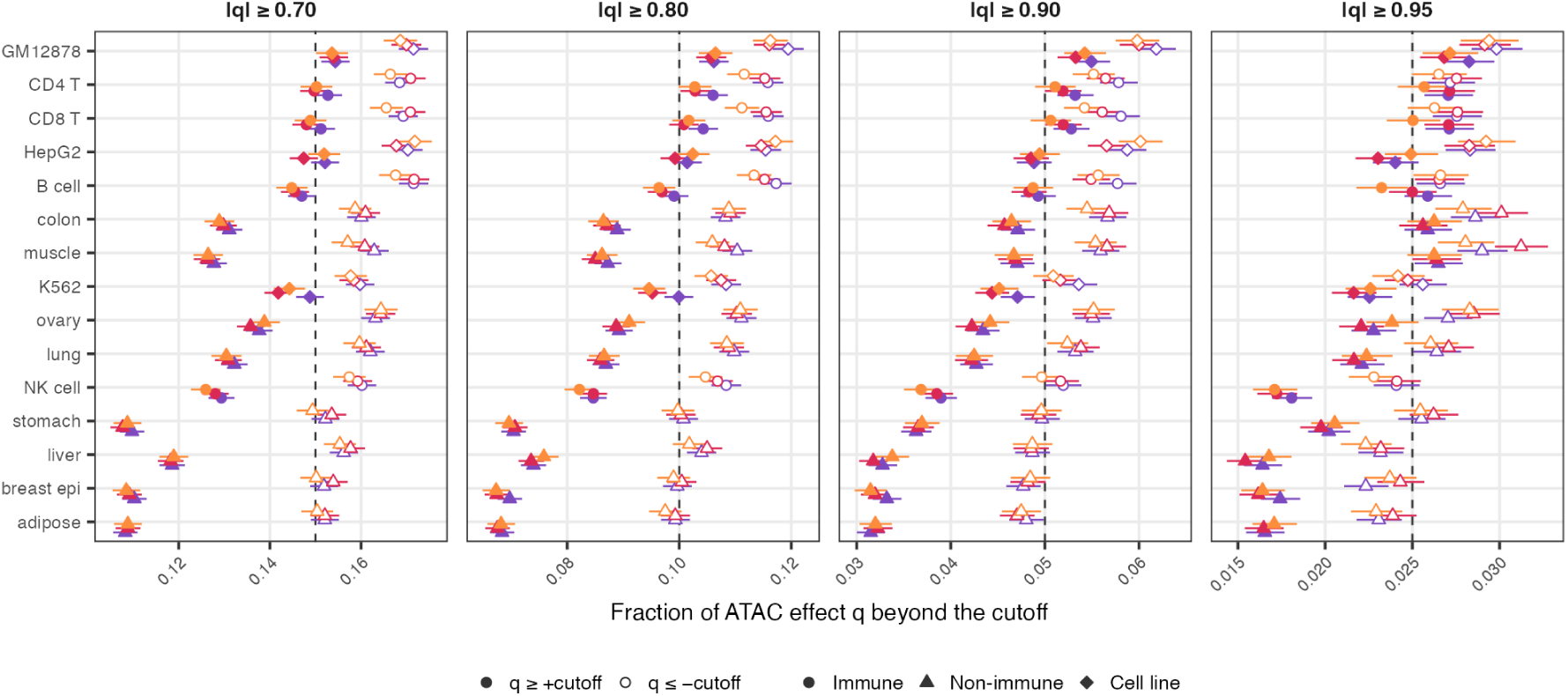
Fraction of introgressed variants beyond the *q* cut-off in each Biosample, at four cut-offs. Filled points are *q ≥* +cut-off and pale points *q ≤ −*cut-off; colour is introgression source and shape is Biosample class. Error bars are exact binomial 95% CIs and the dashed line is the one-sided expectation. Biosample order is fixed to the Denisovan ranking at *|q| ≥* 0.9 in all four panels.

**Supplementary Figure 8.**
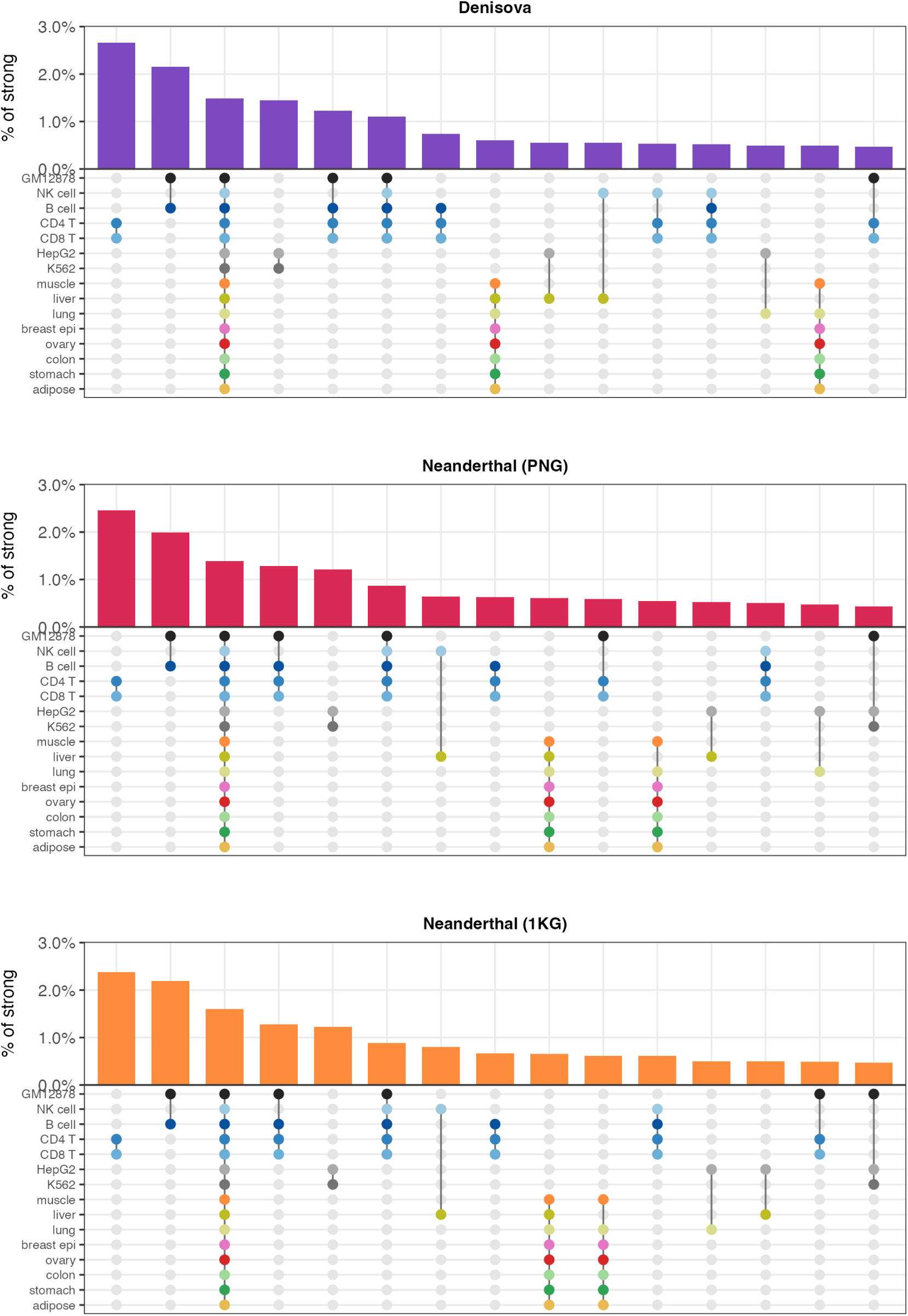
Top 15 most frequent multi-Biosample combinations among variants with *|q| ≥* 0.9 within each introgression source, as a percentage of that source’s set.

**Supplementary Figure 9.**
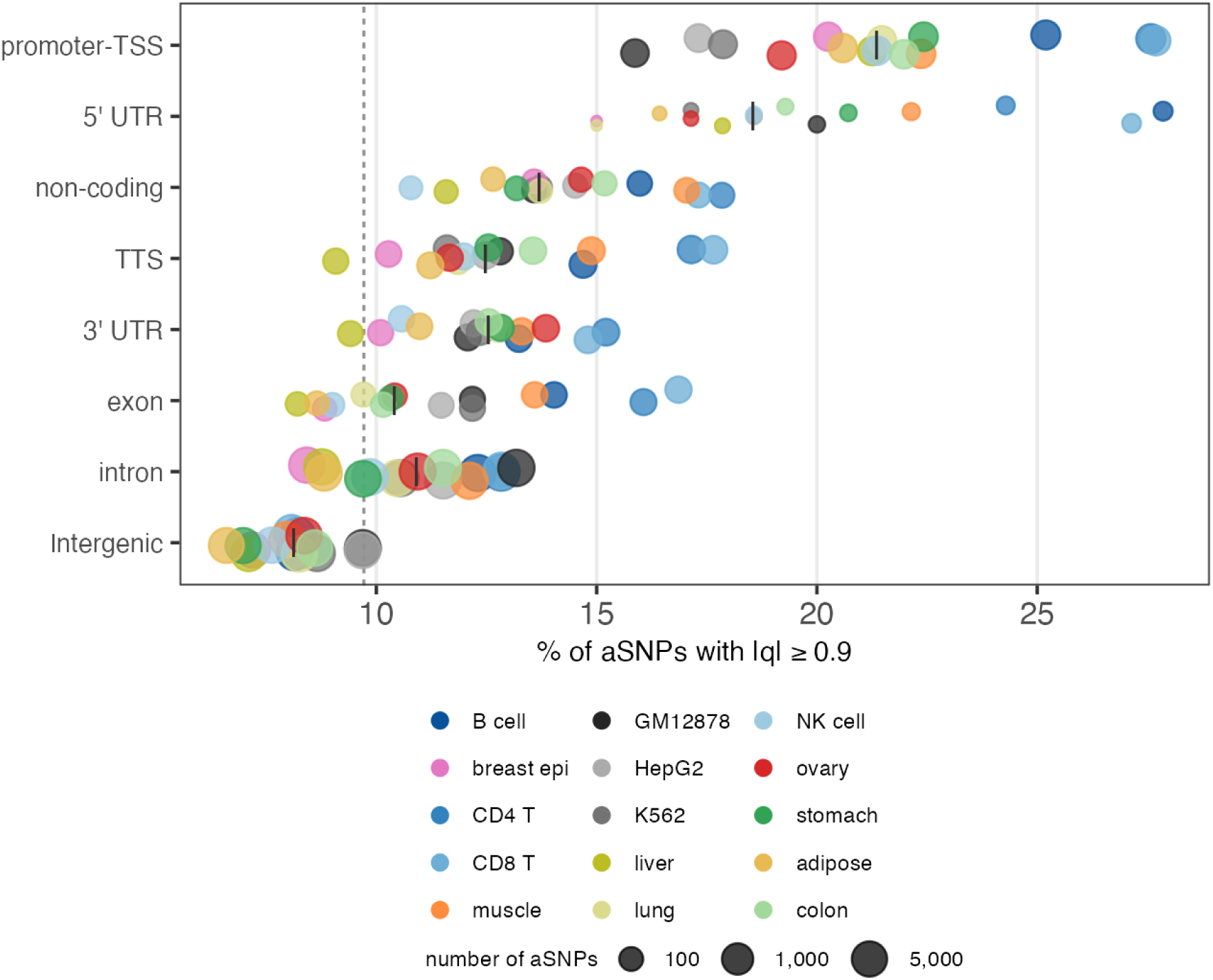
Percentage of introgressed variants reaching *|q| ≥* 0.9 within each HOMER annotation class. The small vertical line in each category indicates the median across Biosamples, and the dashed line the overall rate across all classes. Classes are ordered by their pooled rate.

**Supplementary Figure 10.**
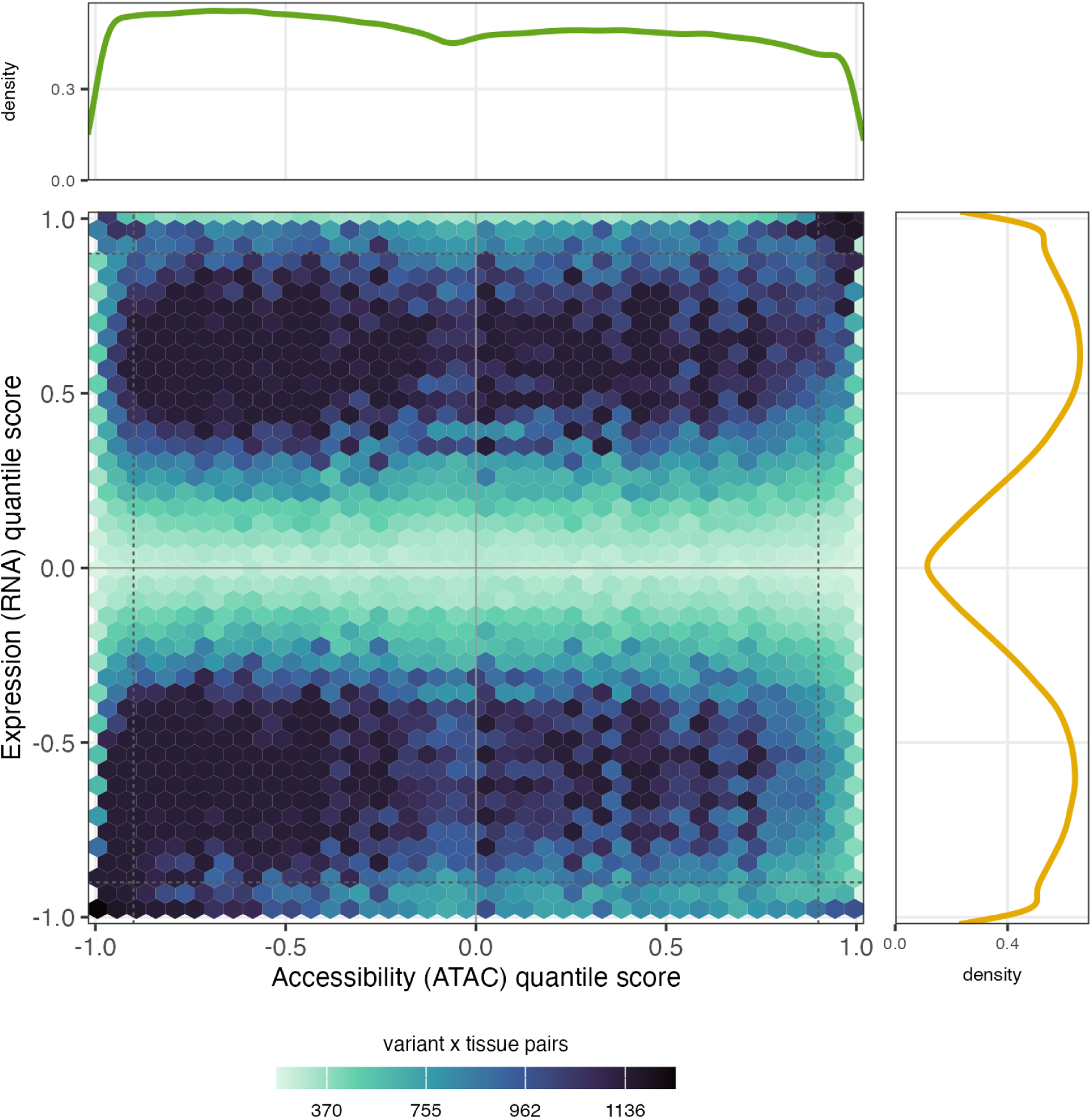
Hexbin plot of AlphaGenome ATAC-seq and RNA-seq effect size quantile score predictions over all variant *×* Biosample pairs scored in both modalities. Bins are coloured according to the number of pairs they contain, and coloured by overall bin density quantile. Dashed lines mark *|q|* = 0.9 in each modality. Marginal panels are the density of each score.

**Supplementary Figure 11.**
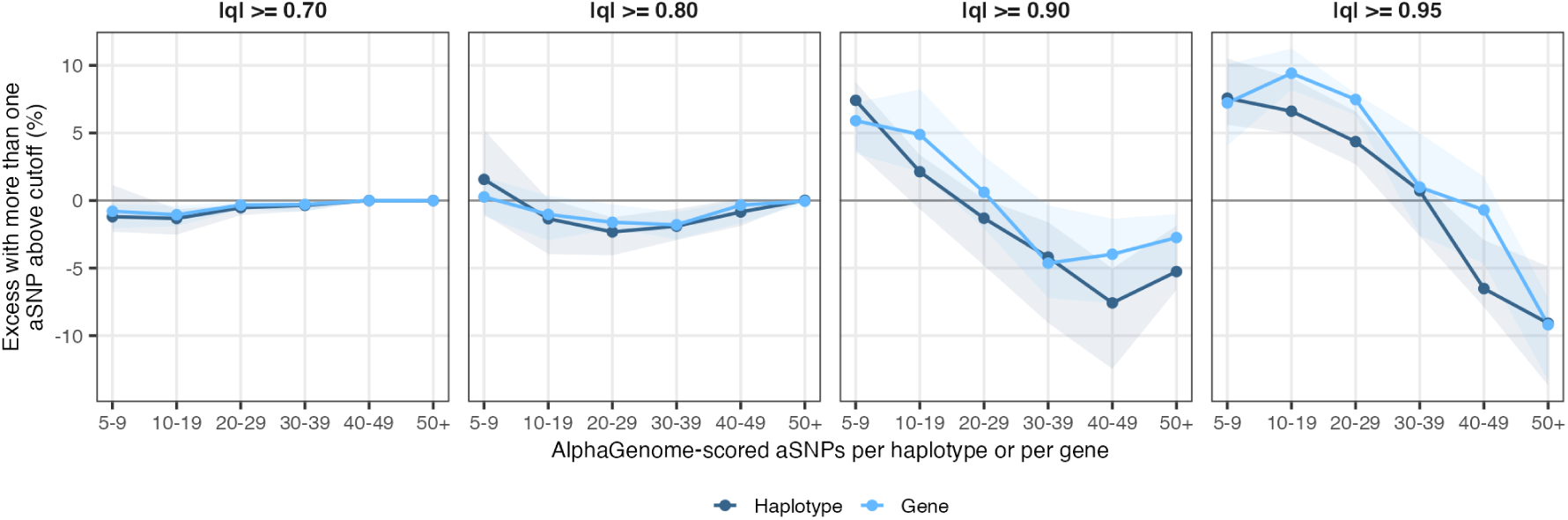
Observed minus expected probability that a haplotype or gene carries more than one aSNP above a *|q|* cut-off, given that it carries at least one, against the number of AlphaGenome-scored aSNPs it contains. The expectation is the Poisson-binomial over each aSNP’s own predicted probability, ribbons are the 10th–90th percentile across the 12 primary tissue Biosamples.

**Supplementary Figure 12.**
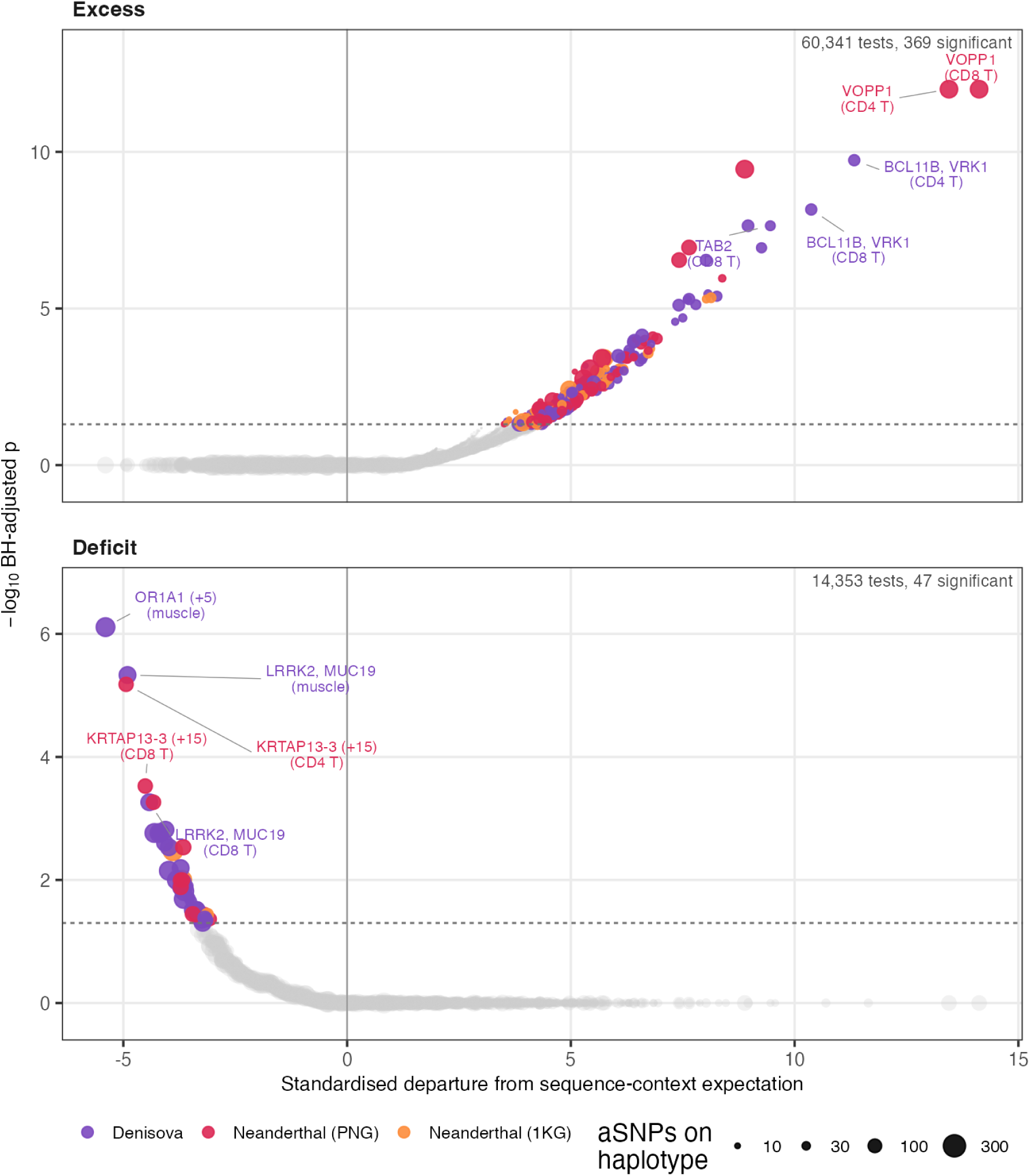
Volcano plot showing departures from expectation against BH-adjusted *p*-value for every haplotype *×* Biosample test at *|q| ≥* 0.9, in each direction. Point area is the number of AlphaGenome-scored aSNPs on the haplotype, colour is introgression source for tests reaching *BHp <* 0.05, and the dashed line is *BHp* = 0.05.

**Supplementary Figure 13.**
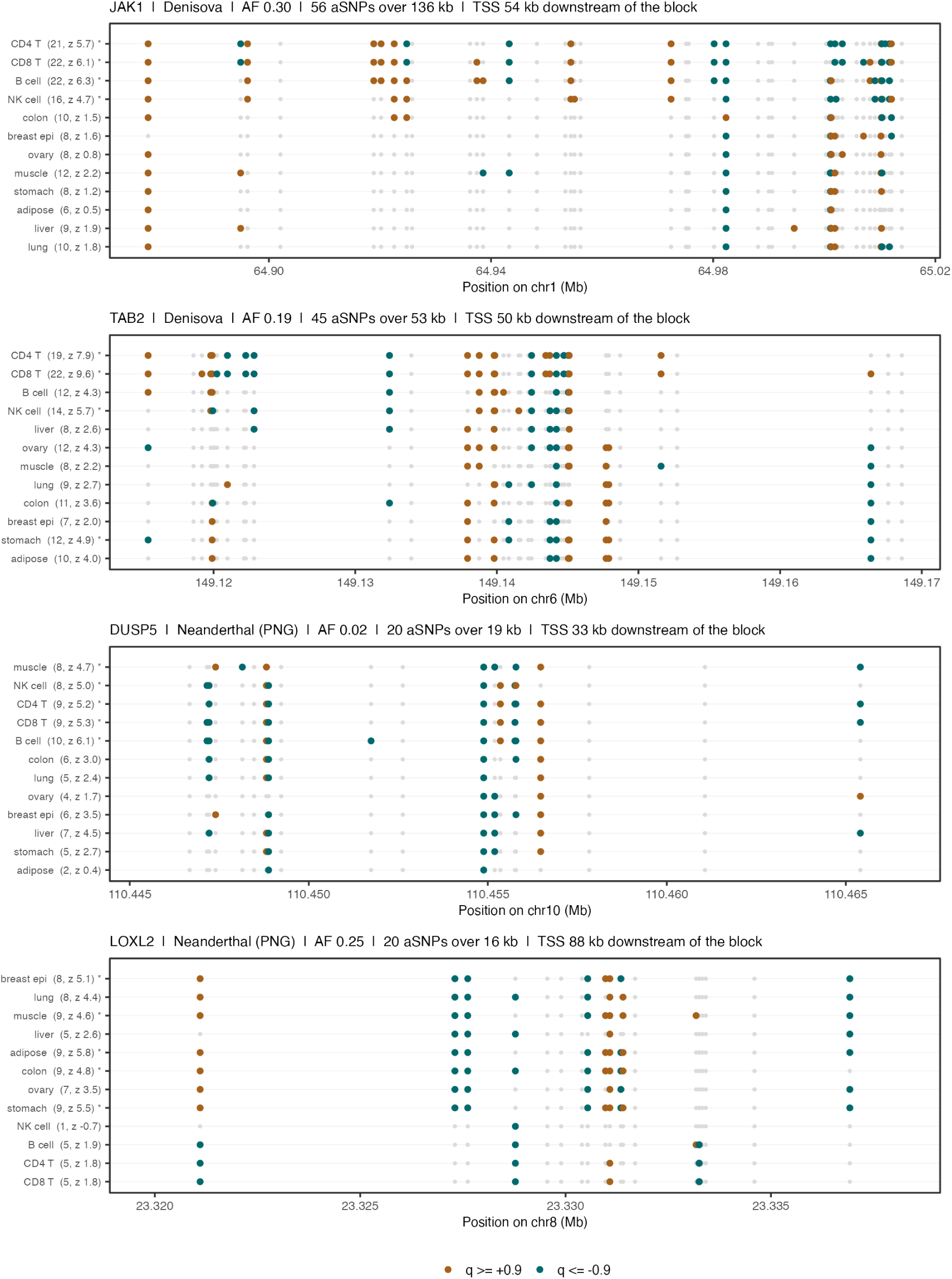
Examples of haplotypes with recurrent excess signal at single-aSNP resolution, one lane per primary Biosample and one point per aSNP at its genomic position. Points are coloured where *q ≥* +0.9 or *q ≤ −*0.9. Numbers in parenthesis indicate the number of aSNPs where *|q| ≥* 0.9 per Biosample, rows with an asterisk are significant after BH correction. Biosamples are ordered by clustering on each haplotype’s own aSNPs. Shaded region indicates the gene body, the vertical line is the TSS.

## Supplementary Tables

**Supplementary table 1**: Biosamples included.

**Supplementary table 2**: Tests for deficit sharing of introgressed haplotypes across biosamples.

**Supplementary table 3**: Tests for excess sharing of introgressed haplotypes across biosamples.

**Supplementary table 4**: Proportion concordant by SNP threshold and dataset (RNA-seq, raw score).

**Supplementary table 5**: Correlation results by TSS distance bin and group, GM12878.

**Supplementary table 6**: *|q| ≥* 0.9 variants by Biosample, with percent shared.

**Supplementary table 7**: GO and Hallmark enrichment among genes near introgressed variants, swept across distance and q-score cutoffs.

**Supplementary table 8**: Significant haplotype-level excess/deficit sharing hits (volcano plot).

